# Real-time neurofeedback to alter interpretations of a naturalistic narrative

**DOI:** 10.1101/2022.01.30.478388

**Authors:** Anne C. Mennen, Samuel A. Nastase, Yaara Yeshurun, Uri Hasson, Kenneth A. Norman

**Affiliations:** Princeton Neuroscience Institute, Princeton University, Princeton, NJ, 08540-1010, USA; School of Psychological Sciences, Tel Aviv University, Tel Aviv 6997801, Israel; Department of Psychology, Princeton University, Princeton, New Jersey 08540-1010, USA

**Keywords:** real-time fMRI, naturalistic stimuli, neurofeedback, shared response model

## Abstract

We explored the potential of using real-time fMRI (rt-fMRI) neurofeedback training to bias interpretations of naturalistic narrative stimuli. Participants were randomly assigned to one of two possible conditions, each corresponding to a different interpretation of an ambiguous spoken story. While participants listened to the story in the scanner, neurofeedback was used to reward neural activity corresponding to the assigned interpretation. After scanning, final interpretations were assessed. While neurofeedback did not change story interpretations on average, participants with higher levels of decoding accuracy during the neurofeedback procedure were more likely to adopt the assigned interpretation. Thus, we believe that individualized neurofeedback shaped interpretations successfully when the signal was accurate, although more work is needed to improve this method and validate the result. While naturalistic stimuli introduce a unique set of challenges in providing effective and individualized neurofeedback, we believe that this technique holds promise for individualized cognitive therapy.

## 1 INTRODUCTION

Compared to healthy participants, depressed participants are more likely to negatively interpret ambiguous scenarios, especially those that contain self-referential prompts (Everaert et al., 2017). Past research has associated increased depression severity with the persistence of these negative interpretations even after learning that the scenarios ended positively (Everaert et al., 2018). To reduce this negative bias and encourage positive interpretations, researchers have used Cognitive Bias Modification for Interpretation (CBM-I) training (Mathews and Mackintosh, 2000). During positive CBM-I training, depressed participants are shown scenarios that differ in interpretation depending on the final word, e.g., *The new people you meet will find you (boring/friendly)*. During training, the final word is revealed to disambiguate the meaning (e.g., *friendly*) so that participants become more likely to expect the positive outcome (Joormann et al.,2015). A meta-analysis collapsing across anxious and depressed individuals found that CBM-I training more effectively reduced negative biases than Attention Bias Modification (ABM) training, which involves training participants to direct their attention away from negatively valenced pictures (Hallion and Ruscio, 2011). One explanation for the increased efficacy is that CBM-I training uses self-relevant stimuli that are inherently more realistic and relatable than the negative faces or isolated words used in ABM training.

As with other training paradigms, however, results are mixed in regards to whether CBM-I training actually improves depressive symptoms (Jones and Sharpe, 2017). One explanation for the mixed clinical results is that the training uses a one-size-fits-all approach. In other words, all participants are trained in the same way, regardless of variability in symptoms, momentary lapses in attention, effort, or belief in one of the interpretations. This raises the prospect that better results could be obtained using individualized training methods with more realistic stimuli. Specifically, given that different interpretations of complex narrative and social scenarios yield different neural responses (Yeshurun et al., 2017), it might be possible to use real-time neuroimaging to track how well participants are adopting the desired interpretation, and to provide feedback to bias them toward the desired interpretation (for recent reviews of real-time fMRI neurofeedback, see Stoeckel et al., 2014; Sitaram et al., 2017; Thibault et al., 2018; Watanabe et al.,2017; Taschereau-Dumouchel et al., 2022; Hampson, 2021).

Before trying this in a clinical setting, we need to demonstrate that it is possible to decode interpretations in real time, and that feedback based on this decoded interpretation is effective in shaping participants’ interpretations. The present study takes some initial steps toward this goal, by assessing whether it is possible to nudge participants’ interpretation of an ambiguous narrative using real-time fMRI (rt-fMRI) neurofeedback.

Here, we build on prior work demonstrating that high-level cortical areas differentiate interpretations of an ambiguous social narrative across individuals (Yeshurun et al., 2017; Finn et al., 2018; Nguyen et al., 2019). Specifically, we based our experiment on a prior study showing that neural responses to an ambiguous 12-minute spoken narrative vary depending on how the story is interpreted (Yeshurun et al., 2017). In this study, Yeshurun et al. (2017) explicitly instructed participants to adopt one of two different interpretations before listening to the story in the fMRI scanner – in one interpretation, the main character’s wife is cheating on him, and in the other interpretation, the main character is just being paranoid (see *Stimulus* section below). This manipulation ensured that the two groups of participants would interpret the story in different ways, allowing the authors to measure neural signatures of the interpretations shared within each group. The authors successfully identified neural regions that accurately predicted the assigned interpretation in held-out participants. Within these regions, the parts of the story that varied most in meaning based on the two interpretations were more likely to yield higher neural classification accuracy.

In the current study, we pre-trained a classifier using the data from Yeshurun et al. (2017) to decode which interpretation participants were adopting; the data from Yeshurun et al. (2017) are openly available as part of the Narratives dataset described in Nastase et al. (2021). Crucially, instead of explicitly instructing participants to adopt a particular interpretation, we randomly assigned participants to an interpretation condition (without telling them which condition they were assigned to) and attempted to nudge them towards the assigned interpretation using neurofeedback. Specifically, while real-time participants listened to the same story in the fMRI scanner, we used the pre-trained classifier to decode which interpretation the participant was most likely thinking about in a given moment, and then provided intermittent neurofeedback to push the participants toward their assigned interpretations - participants were rewarded when the classifier’s estimate of their interpretation of the story matched the participant’s randomly assigned interpretation condition.

After listening to the story, participants answered questions to assess if neurofeedback successfully biased story interpretations. Thus, the critical comparison was between each participant’s actual interpretation at the end of neurofeedback and the target interpretation determined by random group assignment. If neurofeedback was successful, participants in each of the assigned groups would be more likely to adopt the target interpretation of that group, causing interpretations between assigned groups to differ. Overall, we did not reliably bias participants toward the target interpretations. However, taking into account the decoding accuracy of the classifier for each participant, we found that participants with the highest decoding accuracy were more likely to choose the target interpretation.

## 2 METHODS

### 2.1 Participants

Twenty-two participants from Princeton University and the surrounding local community consented to participate in this study, but 2 participants did not return for their second visit. Thus, 20 participants were included in the analysis (12 female, 2 left-handed, mean age = 20.5 years). Participants received monetary compensation for their participation, including an additional bonus based on their neurofeedback performance ($20 maximum). The study was approved by Princeton University’s Institutional Review Board.

### 2.2 Stimuli

All participants listened to a 12-minute adapted version of “Pretty Mouth and Green My Eyes” by J. D. Salinger, read by a professional actor. The same recording was used in Yeshurun et al. (2017). The audio stimulus began with 18 s of music and ended in silence; see Yeshurun et al. (2017) for further details about the stimulus. The audio stimulus is openly available as part of the Narratives dataset (Nastase et al., 2021).

The story begins when the character Arthur calls his friend, Lee, after returning home late at night without his wife, Joanie. Arthur is concerned about Joanie’s whereabouts. Lee is at home in bed with a woman, although the narrator purposefully leaves her identity ambiguous. Yeshurun et al. (2017) created two interpretation groups by briefing participants on this woman’s identity *before* they heard the story. The two interpretation groups were:

1. **cheating**: Joanie is the woman in Lee’s bed. In this case, Joanie is cheating on her husband, Arthur, with Lee.
2. **paranoid**: Lee’s girlfriend, Rose, is the woman in Lee’s bed. In this case, Arthur is paranoid that Joanie is cheating on him and is bothering Lee late at night.

### 2.3 Procedure

On the first visit, participants consented to the experiment and underwent initial structural and functional scans for image registration. Because our pre-trained classifier was trained in standard MNI space, registering each participant’s brain to MNI space before Visit 2 allowed us to apply the classifier in real-time. For registration details, see Methods, Sections 2.4.1 and 2.4.2.

On the second visit (at least one day later, mean delay = 4.6 days), participants returned to complete 4 runs of neurofeedback training (one participant only completed 2 runs due to technical problems). Before training began, participants were randomly assigned in a double-blind fashion into either the cheating or paranoid interpretation group. Group assignment was performed via a Python script that saved the assignment as a text file to be used later during neurofeedback. This way, both the experimenter and participant were blind to group assignment. Of the 20 participants, 10 were randomly assigned to the cheating group and 10 were randomly assigned to the paranoid group. Importantly, the assigned interpretation group for a particular participant was the same across all 4 runs of neurofeedback training (i.e., if a participant was assigned to the cheating group for the first run, they were also in the cheating group for runs 2, 3, and 4).

As noted above, we did not tell participants which interpretation to take at the onset of the experiment. Instead, we informed them of the two possible interpretations and that both were equally likely (meaning that there was no ground truth or correct interpretation). Thus, they would have to use neurofeedback to determine which was the correct interpretation for them.

While participants listened to the story, we used a visual stimulus to present neurofeedback (Figure 1). For most of the story, participants viewed a gray rectangle, indicating that they should simply continue listening. Four seconds prior to a specific period in the story when we would analyze brain activity (which we refer to as a *station*), participants indicated which interpretive “lens” (cheating or paranoid) they were going to adopt for the upcoming station. Participants were free to choose either interpretation at any time, as long as they were trying to maximize their scores. Participants pressed their index or middle fingers to indicate their choice of either the cheating or paranoid interpretation (left/right position of choices was counterbalanced across participants).

**FIGURE 1.**
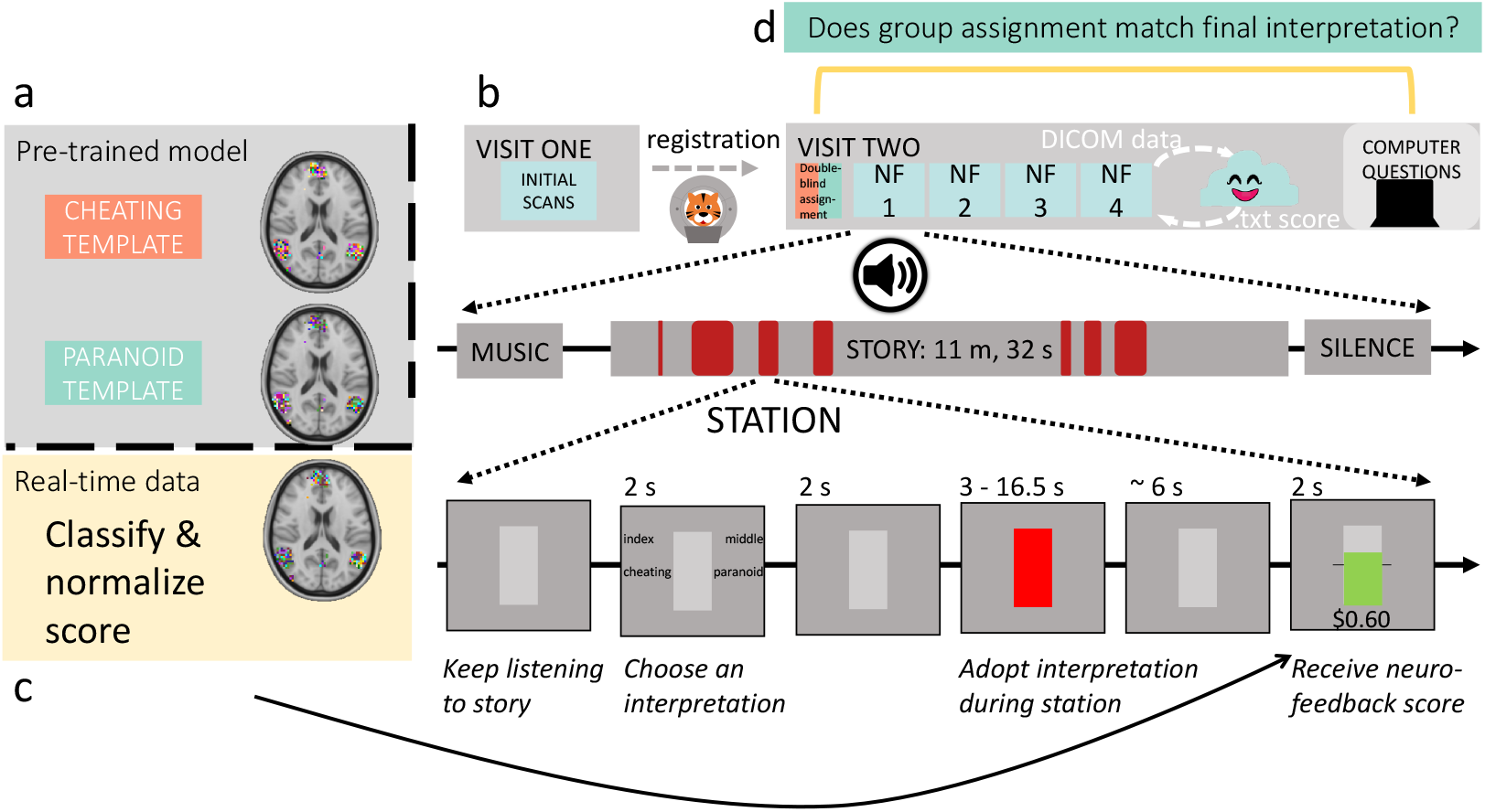
Experimental design. **(a)** Before scanning, we pre-trained a classification model on data from Yeshurun et al. (2017) to create template brain responses at each station. **(b)** On Visit 1, a high-resolution anatomical scan and a short functional scan were acquired for registration to MNI space. At least one day later, participants returned for 4 runs of story listening with neurofeedback training. Each run was about 12 minutes long, and featured the same audio stimulus used by Yeshurun et al. (2017). Each run included 7 *stations* where brain activity was analyzed. Stations varied in length between 3 - 16.5 seconds. **(c)** After each station, the neurofeedback score was displayed and participants received a reward based on the model prediction for that station, normalized by pilot experimental data. **(d)** Afterward, participants answered questions outside of the scanner to assess their final interpretations of the story.

Two seconds after the probe, the station began. During these stations, the rectangle turned red to signal that brain activity was being recorded. We did this so that participants were aware of when we were analyzing their brain activity. We explicitly marked stations to help participants determine which thoughts led to higher rewards, in hopes of decreasing the credit-assignment problem inherent in the delayed rt-fMRI signal (Oblak et al., 2017); see Appendix A.3 for details on how we chose the stations. To account for hemodynamic lag, the BOLD data that we analyzed for each station were shifted by 3 TRs (4.5 s) from the actual TRs when the “station recording signal” was on screen.

After each station, the data were preprocessed and the pre-trained model was used to decode the participant’s interpretation at that station. The rectangle then displayed the neurofeedback score; a horizontal line through the center indicated the threshold to earn any extra money and the filled area corresponded to the score. If participants scored above threshold, the score was shown in green with the monetary reward underneath. Otherwise, the score was shown in gray with $0.00 below the rectangle. Because of the aforementioned 3-TR hemodynamic shift, plus one TR of time for data analysis, participants typically had to wait 4 TRs (6 s) for the feedback display to appear after the end of the station (see Appendix B for more details regarding feedback timing).

To assess the final interpretations after all runs, participants answered the same questions from Yeshurun et al. (2017). The 39 questions contained 27 comprehension questions (e.g., *What was the girl doing when the phone rang?*) and 12 questions that directly probed interpretation (e.g., *Did you think Joanie was cheating on Arthur?*). Additionally, participants provided numerical ratings (spanning 1-5) indicating their opinions on various topics, such as how much they empathized with the characters, enjoyed the story, thought neurofeedback helped, etc. Lastly, participants completed a survey to assess perception and strategies.

*Note*: We emphasized the following in our instructions before scanning began, to make sure that participants were attending to neurofeedback throughout the entire experiment:

- The story was written to be purposefully ambiguous, instead of there being one true interpretation.
- You should try to interpret the story based on your neurofeedback, not what you think was intended by the author.
- Even if you think that the correct interpretation was revealed in the story, characters might be lying. Keep using neurofeedback to guide your interpretations.
- The neurofeedback scores reflect how clearly and correctly your brain is interpreting the story. Low scores can indicate either (1) noisy signals (e.g., poor focus) and/or (2) incorrect interpretations.
- The neurofeedback score only represents the neural information recorded during the station. We keep track of your responses to the probes for analysis purposes, but the neurofeedback score you see is only based on your brain activity.

The full set of instructions used in the experiment is included in Appendix D, Figure 16.

After scanning, participants completed a questionnaire and survey. Figure 1 illustrates the full experimental design and the neurofeedback stimuli.

### 2.4 Data acquisition

All scanning data were acquired with a 3T MRI scanner (Siemens Skyra) and a 64-channel head coil. Sequences were matched to Yeshurun et al. (2017) as closely as possible. Both scanning sessions began with a Siemens scout scan for automated slice alignment to the ACPC axis. On Visit 1 only, we collected a high-resolution, T1-weighted magnetization-prepared rapid acquisition gradient-echo (MPRAGE) anatomical scan to facilitate normalizing each participant’s functional data to standard space: repetition time (TR) = 2,300 ms, TE = 3.08 ms, flip angle = 9°, resolution = 0.86 x 0.86 x 0.9 mm^3^, FOV = 220 mm^2^.

We ran functional scans on Visit 1 for registration purposes only (i.e., without presenting stimuli); and we ran functional scans for the experiment during Visit 2. All functional scans used a T2*-weighted echo-planar imaging sequence: 1.5 s TR, 28 ms echo time, flip angle =64°, 3 x 3 x 4 mm^3^ voxel size, 64 x 64 matrix, 192 x 192 mm^2^ field of view, 27 slices, no gap between slices, interleaved slice acquisition.

No fieldmap scans were collected, as we were trying to best match the real-time data to the data used to estimate the pre-trained model. As the data from Yeshurun et al. (2017) did not include fieldmap scans, we omitted real-time susceptibility distortion correction in this experiment.

#### 2.4.1 Offline image registration

The following sections describe how we registered each participant to MNI space after the Visit 1 scans, in preparation for Visit 2. We used *fMRIPprep* 1.2.3 (Esteban et al., 2019; Esteban et al., 2018; RRID:SCR_016216), which is based on *Nipype* 1.1.6-dev (Gorgolewski et al., 2011; Gorgolewski et al., 2018; RRID:SCR_002502). The following description of preprocessing was generated with fMRIPrep.

The T1-weighted (T1w) image was corrected for intensity non-uniformity (INU) using `N4BiasFieldCorrection` (ANTs 2.2.0; Tustison et al., 2010), and used as T1w-reference throughout the workfìow. The T1w-reference was then skull-stripped using `antsBrainExtraction.sh` (ANTs 2.2.0), using OASIS as target template. Brain surfaces were reconstructed using `recon-all` (FreeSurfer 6.0.1, RRID:SCR_001847, Dale et al., 1999), and the brain mask estimated previously was refined with a custom variation of the method to reconcile ANTs-derived and FreeSurfer-derived segmentations of the cortical gray-matter of Mindboggle (RRID:SCR_002438, Klein et al., 2017). Spatial normalization to the ICBM 152 Nonlinear Asymmetrical template version 2009c (Fonov et al., 2009, RRID:SCR_008796) was performed through nonlinear registration with `antsRegistration` (ANTs 2.2.0, RRID:SCR_004757, Avants et al., 2008), using brain-extracted versions of both T1w volume and template. Brain tissue segmentation of CSF, WM and GM was performed on the brain-extracted T1w using fast (FSL 5.0.9, RRID:SCR_002823, Zhang et al., 2001).

For the BOLD run collected during Visit 1, the following registration processing was performed. First, a reference volume and its skull-stripped version were generated using a custom methodology of *fMRIPrep*. The BOLD reference was then co-registered to the T1w reference using `bbregister` (FreeSurfer) which implements boundary-based registration (Greve and Fischl, 2009). Co-registration was configured with nine degrees of freedom to account for distortions remaining in the BOLD reference. Head-motion parameters with respect to the BOLD reference (transformation matrices, and six corresponding rotation and translation parameters) were estimated before any spatiotemporal filtering using `mcflirt` (FSL 5.0.9, Jenkinson et al., 2002). The BOLD time series was resampled onto the original, native space by applying a single, composite transform to correct for head-motion. The BOLD time series was then resampled to MNI152NLin2009cAsym standard space. First, a reference volume and its skull-stripped version were generated using a custom methodology of *fMRIPrep*. All resamplings were performed with *a single interpolation step* by composing all the pertinent transformations (i.e. head-motion transform matrices and co-registrations to anatomical and template spaces). Gridded (volumetric) resamplings were performed using `antsApplyTransforms` (ANTs), configured with Lanczos interpolation to minimize the smoothing effects of other kernels (Lanczos, 1964).

Many internal operations of *fMRIPrep* use *Nilearn* 0.4.2 (Abraham et al., 2014, RRID:SCR_001362), mostly within the functional processing workflow. For more details of the pipeline, see the section corresponding to workflows in fMRIPrep’s documentation.

#### 2.4.2 Real-time image registration and processing

As BOLD data arrived from the scanner, the data were transferred as bytes to a secure cloud server, where all subsequent processing steps were performed. The data were returned to the local Linux as a .txt file containing the final neurofeedback score to be displayed. Real-time processing was handled using the RT-Cloud software package (Wallace et al., 2022; Kumar et al., 2020); see Appendix B for full details on the cloud setup.

BOLD data were acquired in the participant’s native space, but the pre-trained model was in standard MNI space. Therefore, we had to transform each incoming BOLD volume to MNI space in real-time. To do this, we combined Visit 1’s previously calculated registration steps from fMRIPrep with real-time registration of each new BOLD volume. In real-time, we used `mcflirt` (FSL 5.0.9, Jenkinson et al., 2002) to register each incoming BOLD volume with the example functional image acquired on Visit 1. We combined this transformation matrix with the two transformation matrices calculated previously with fMRIPrep: (1) the transformation from functional → T1w space, and (2) the transformation from T1w space → MNI space. All transformations were concatenated and performed in a single step using `antsApplyTransforms` (Avants et al., 2008). Details of this pipeline performed on the cloud server are shown in Table 2 in Appendix B.

### 2.5 Classification

We used 7 pre-trained logistic regression classifiers (sklearn solver = ‘lbfgs’, C = 1), corresponding to the 7 stations. See Appendix A for details on classifier construction. Each classifier estimated *p*(*c*), the probability that the participant was interpreting the story in line with the cheating interpretation (the probability of the paranoid interpretation was 1 - *p*(*c*)). We generated this probability estimate using scikit-learn’s predict_proba function (Pedregosa et al., 2011). For more detailed information on how we optimized the classifier design, see Appendix A.

To convert this prediction to the neurofeedback score delivered to the participant, we normalized *p*(*c*) based on pilot data. We learned from the pilot experiment (Appendix C) that - in our paradigm, where participants are allowed to form their own interpretations, as opposed to being explicitly told which interpretation to use as in Yeshurun et al. (2017) — the mean *p*(*c*) values varied considerably across stations, such that many stations were strongly biased toward particular interpretations (regardless of group assignment). To control for these biases, we decided to provide neurofeedback based on participants’ deviation from the “average neural interpretation trajectory” (where this “average neural interpretation trajectory” was computed by collapsing results across the two conditions in the pilot study), rather than providing neurofeedback based on whether participants’ decoded interpretation matched the assigned interpretation at a particular moment. That is, at a moment when the narrative leans toward the paranoid interpretation on average, we decided to reward participants in the cheating group if their interpretation was closer to cheating than the average, even if (in absolute terms) their interpretation was closer to paranoid than to cheating. This kind of neurofeedback can be viewed as “nudging” individual trajectories off the default path.

Specifically, for each participant, at a given station *st*, we applied the following formula to transform *p*(*c*) to a neurofeedback score:

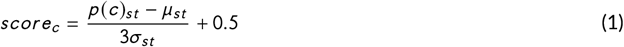

where *μ_st_* and *σ_st_* were the mean and standard deviation, respectively, of *p*(*c*) for that station, from all participants and runs in the pilot experiment. We included a scaling factor of 3 so that the z-scored differences would range roughly between [-0.5,0.5]. Thus, by adding 0.5 to this ratio, we could set the range to be [0,1]. In case scores were much larger or much lower than station means, we added the thresholds:

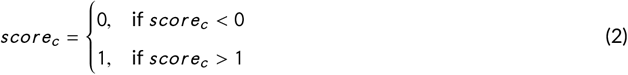

Next, we generated a score based on group assignment with:

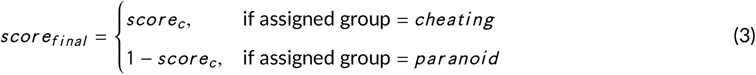

In this framework, participants received a neurofeedback score of .5 if their *p*(*c*) was equal to the mean across participants in the pilot experiment. To earn a reward, participants had to score *above* the station’s mean in the assigned direction. To enforce this constraint, neurofeedback scores ≤ .*5* were converted to 0 values:

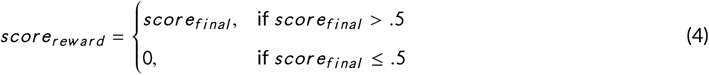

In the results that follow, when reporting neurofeedback scores, we report this final value where scores ≤ 0.5 were converted to 0, since this corresponds to the actual rewards received by participants.

### 2.6 Story comprehension and interpretation scores

Participants answered 39 story questions: 27 general comprehension-based questions and 12 interpretation-specific questions (Yeshurun et al., 2017). We did not analyze one of the interpretation-specific questions that asked for Lee’s girlfriend’s name, since we informed participants that Lee had a girlfriend named Rose prior to the experiment. All participants answered this question correctly, regardless of their interpretation.

To score the interpretation-specific questions, we assigned a +1 score to each answer corresponding to the cheating interpretation, and a −1 score to each answer corresponding to the paranoid interpretation. To calculate a final score, we totaled all interpretation-specific scores and divided by the total number of questions. Thus, a +1 indicates answering all questions consistent with a cheating interpretation, and a −1 indicates answering all questions consistent with a paranoid interpretation.

To collapse across assigned groups and compare “correctness” of final interpretations (i.e., consistency with group assignment), we transformed the interpretation-specific scores by multiplying the scores for participants assigned to the paranoid group by −1. After this transformation, a +1 score indicates answering all questions consistently with one’s group assignment (correct), and a −1 score indicates answering all questions inconsistently with one’s group assignment (incorrect). When we use these transformed scores in the analyses below, we refer to them as “correct interpretation” scores.

### 2.7 Empathy ratings

After answering the story questions, participants rated how much they empathized with Arthur, Lee, Joanie, and “the girl” (the mysterious woman in Lee’s bed). We expected those who assumed the cheating interpretation to empathize with Arthur, and not with Lee and Joanie. Likewise, we expected those who believed the paranoid interpretation to feel more empathy for Lee and Joanie, instead of Arthur. To compute one score that captured the empathy bias, we calculated the difference between each participant’s empathy ratings for Arthur and Lee. Analogously to how we computed “correct interpretation” scores, we also computed “correct empathy” scores by multiplying the empathy difference for all participants assigned to the paranoid group by −1. Thus, positive “correct empathy” scores indicate that the participant empathized with Arthur and Lee in a manner that was consistent with the assigned interpretation group.

## 3 RESULTS

### 3.1 Story comprehension and interpretation scores

Comprehension scores indicated that all participants understood the story (Figure 2a); there was no significant difference between assigned groups (t(18) = 1.41, p = 0.17). Our neurofeedback manipulation did not push participants to their assigned interpretation, as indicated by the interpretation scores. Figure 2b shows the interpretation scores by assigned group. Neither group showed significant bias toward their assigned interpretation (all p > 0.20). Further, the groups did not differ in interpretation scores (one-tailed t(18) = −0.062, p = 0.48).

**FIGURE 2.**
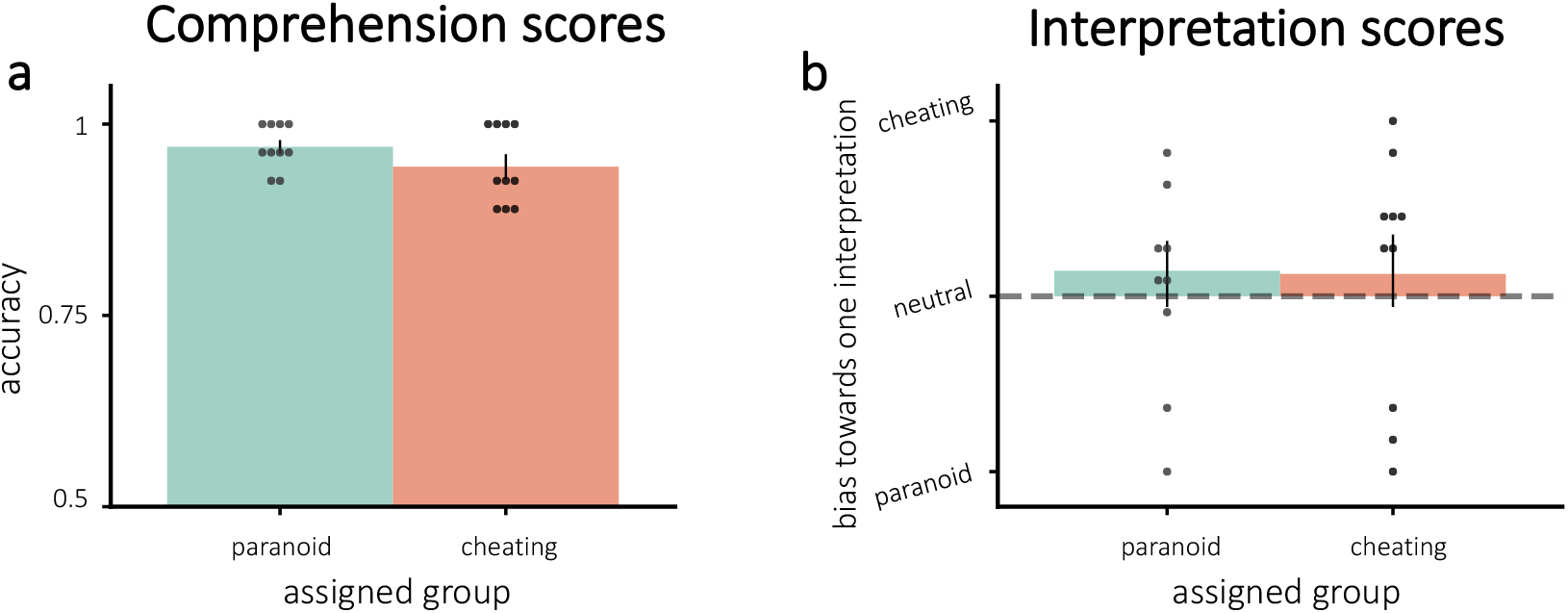
Average scores for **(a)** comprehension and **(b)** interpretation questions. **(a)**: All participants understood the story. **(b)**: Interpretations were not modified by neurofeedback - neither group was significantly pushed toward their assigned interpretation. Error bars = ±1 s.e.m.

### 3.2 Empathy ratings

The empathy ratings for some of the characters differed by group in the “correct” direction of the assigned interpretations. There was a nonsignificant trend for participants in the cheating group to have more empathy for Arthur than participants in the paranoid group (one-tailed t(18) = 1.38, p = 0.093). Additionally, participants in the paranoid group had more empathy for Lee than the participants in the cheating group (one-tailed t(18) = −1.76, p = 0.048). The difference in empathy for Arthur and Lee was significantly different between the assigned groups, with those assigned to the cheating condition having more empathy for Arthur than Lee (one-tailed t(18) = 1.84, p = 0.041). Figure 3 plots the empathy ratings by assigned group. Additionally, there was a significant positive correlation between the difference in empathy for Arthur and Lee and interpretation scores (Pearson r = 0.47, p = 0.037), implying that the more participants empathized with Arthur over Lee, the more likely they were to endorse the cheating interpretation in response to the interpretation questions.

**FIGURE 3.**
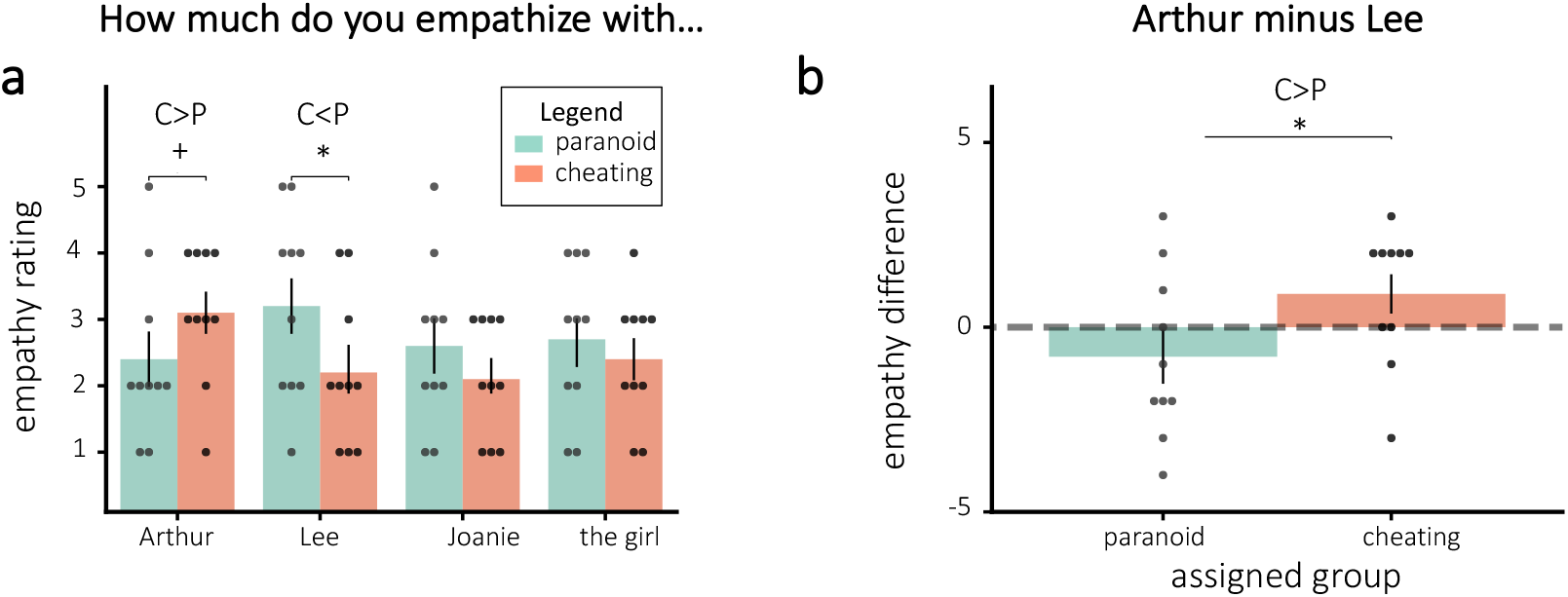
Empathy ratings for **(a)** all characters and **(b)** the difference for Arthur and Lee, separated by assigned group. **(a)**: Empathy ratings differed significantly in the predicted direction for Lee based on group assignment. **(b)**: The difference in empathy for Arthur and Lee also differed significantly by assigned group. Error bars = ±1 s.e.m. * = p < 0.05; + = p < 0.1

### 3.3 Neurofeedback scores

If neurofeedback successfully modified neural responses to the story, we would expect participants to differ in the neurally-decoded cheating probability based on random group assignment. Figure 4a plots cheating probability *p*(*c*) across all stations and runs. Against our expectations, participants in the cheating group did not have significantly larger *p*(*c*) during any individual runs - in fact, *p*(*c*) was numerically smaller for participants in the cheating group (compared to the paranoid group) in the fourth run. Similarly, participants did not show a significant improvement in neurofeedback scores from run 1 to run 4 (combining results from both groups, neurofeedback scores numerically decreased on average; neurofeedback scores numerically increased for the paranoid group and numerically decreased for the cheating group; Figure 4d). Thus, we were unable to reliably push neurofeedback scores, both in terms of *p*(*c*) and normalized neurofeedback rewards, in the assigned group direction.

**FIGURE 4.**
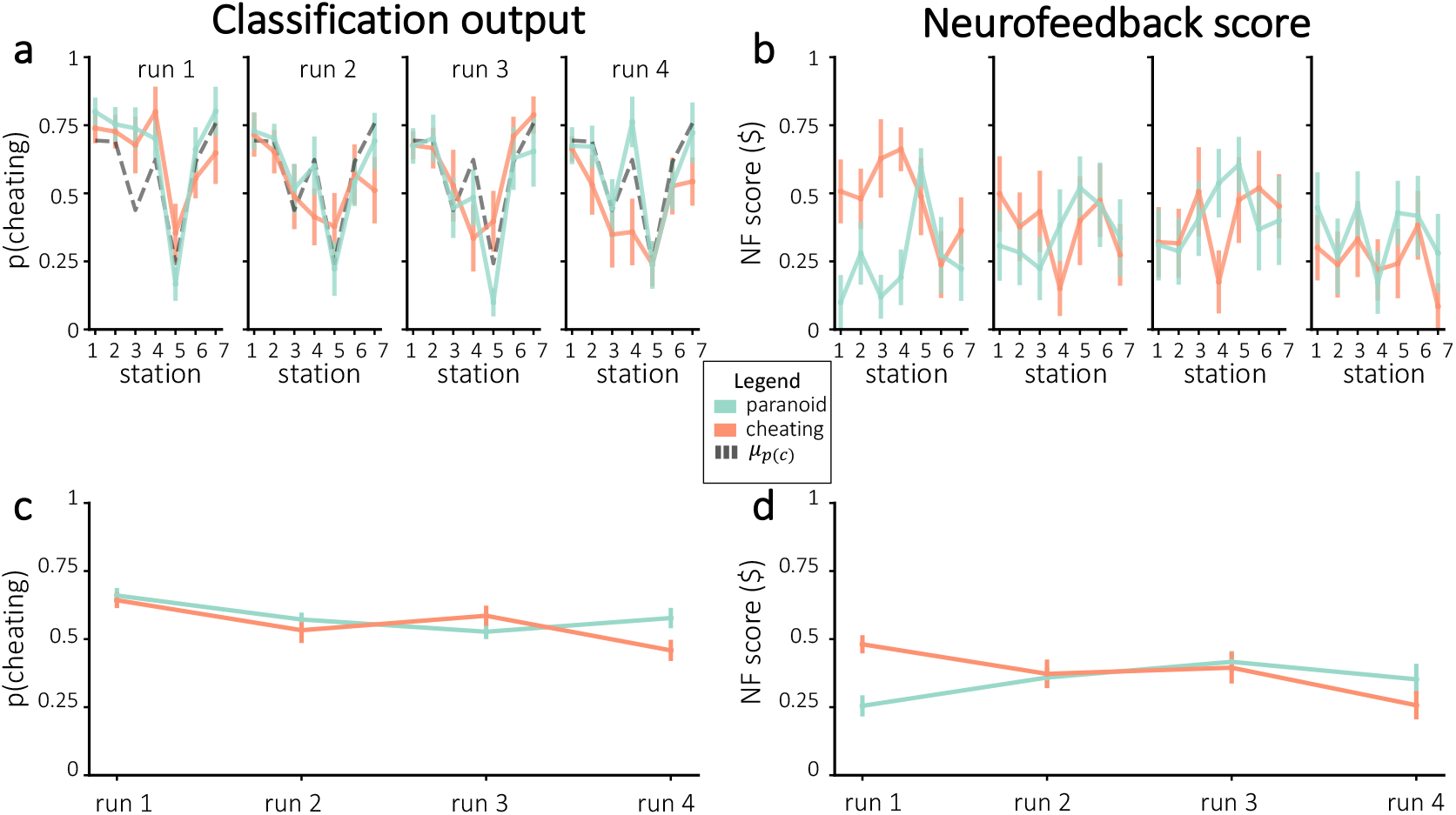
Neurofeedback results, in terms of *p*(*c*) **(a, c)** and normalized neurofeedback score **(b, d)**. **(a)** Average cheating probability (not normalized), separated by assigned group. The dashed line represents mean *p*(*c*) from the pilot experiment. **(b)** Neurofeedback scores that were delivered to participants during real-time neurofeedback, divided by assigned group. Figures **(c-d)** show the run-wise averages for the same values shown in **(a-b)**, respectively. Error bars = ±1 s.e.m.

### 3.4 Probe responses

Prior to each station, we probed participants as to which interpretation they were adopting for this station. Figure 5 shows the behavioral probe responses that were chosen, divided by assigned interpretation groups. While the average choices numerically diverged between groups in the assigned or “correct” direction for each of the runs, none of these run-wise differences were significant after correction for multiple comparisons.

**FIGURE 5.**
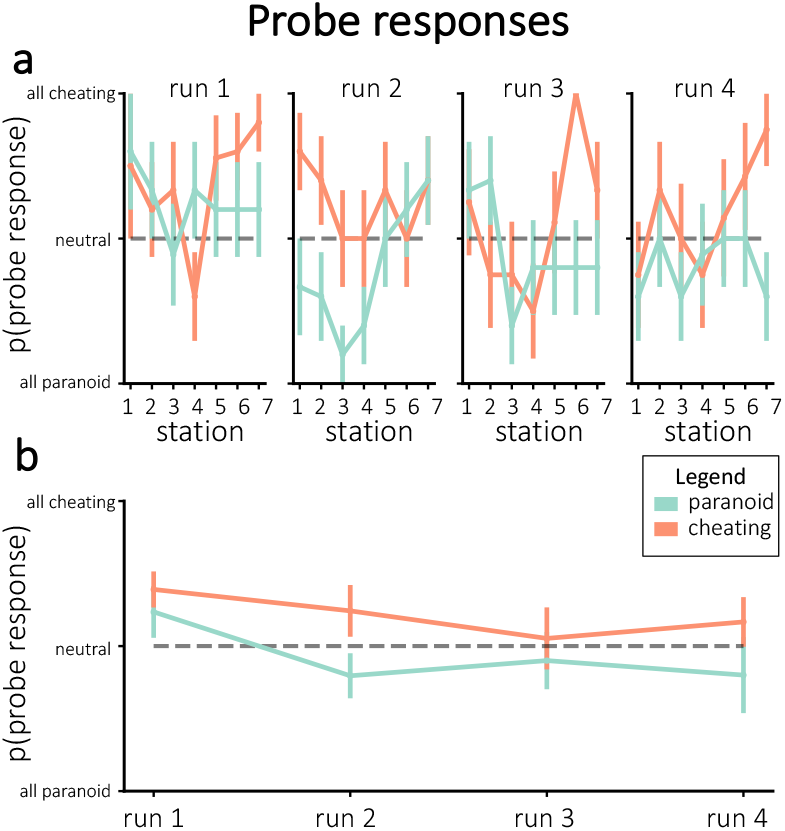
**(a)** Average probe responses during neurofeedback for individual stations, separated by assigned group. **(b)** Average probe responses during neurofeedback, collapsed across stations within each run and separated by assigned group. None of the run-wise differences were significant after correcting for multiple comparisons. Error bars = ±1 s.e.m.

### 3.5 Decoding accuracy

By collecting behavioral probe responses at each station, we were able to measure how accurately the classifier was decoding story interpretations in each participant. If 1) a participant was truly adopting distinct interpretations when they gave a probe response of “cheating” vs. “paranoid”, and 2) the classifier was sensitive to these interpretive differences, then we would expect to see higher values of *p*(*c*) (i.e., the classifier’s estimate of the strength of the cheating interpretation) for stations where participants (behaviorally) reported thinking of the cheating interpretation compared to the paranoid interpretation. To assess this, for each participant, we separated all classification probabilities by probe response. We averaged *p*(*c*) over all of the times the participant gave a “cheating” probe response and (separately) over all of the times the participant gave a “paranoid” probe response. We computed *decoding accuracy* for each participant by taking the difference of the average *p*(*c*) value for “cheating” probe responses and the average *p*(*c*) value for “paranoid” probe responses. Note that low decoding accuracy can have multiple causes - the participant might not be adopting distinct interpretations, or the classifier might be insensitive (see *Discussion*) - but high decoding accuracy indicates that, to some degree, the participant is adopting distinct interpretations and the classifier is detecting them.

On average, there was a nonsignificant trend for participants to show above-zero decoding accuracy (i.e., larger *p*(*c*) values for stations where participants endorsed the cheating interpretation compared to the paranoid interpretation), one-tailed t(19) = 1.40, p = 0.088. Results are shown in Figure 6a. The figure also shows that there was considerable variation across participants in the level of decoding accuracy.

**FIGURE 6.**
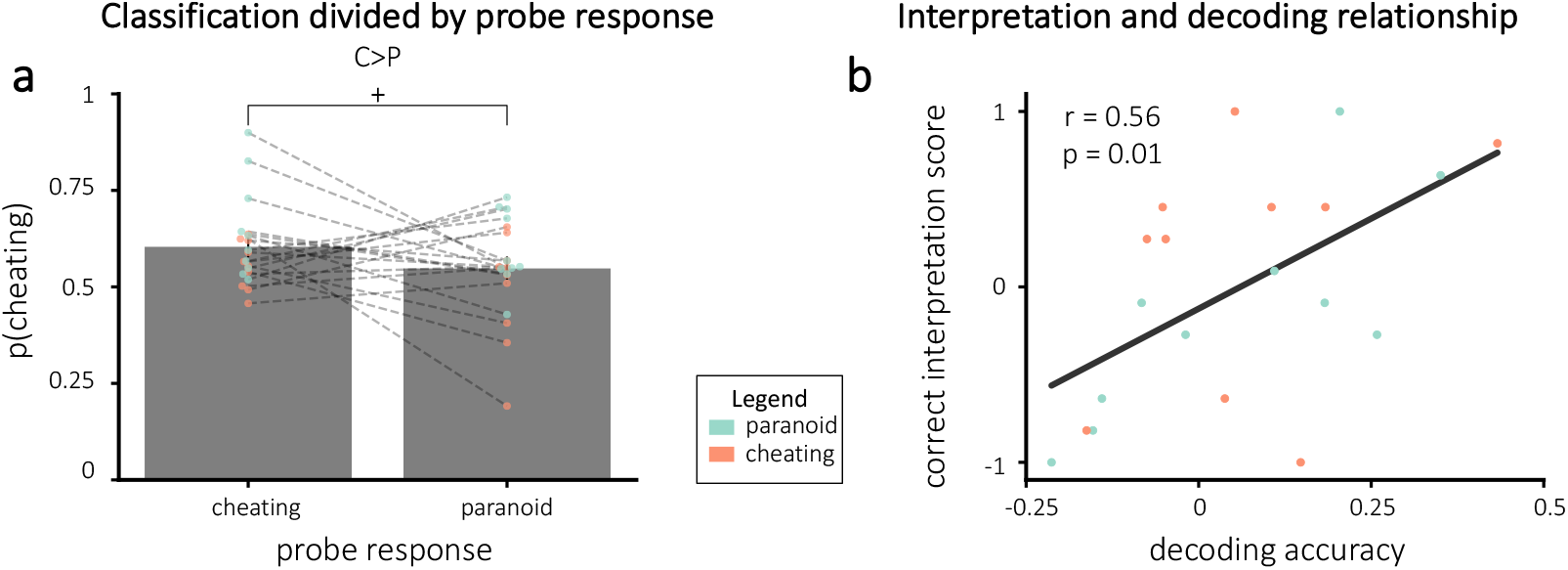
Decoding accuracy results overall **(a)** and related to correct interpretation scores **(b)**. **(a)** Cheating probability was numerically higher when participants endorsed the cheating interpretation compared to the paranoid interpretation, but this difference did not reach statistical significance. *Note*: the x-axis represents the probe responses at each station, *not* the assigned interpretation group. Lines connect each participant’s average probability for each response. **(b)** Correct interpretation plotted as a function of decoding accuracy. Decoding accuracy and correct interpretation scores were positively correlated, suggesting that participants with accurate neurofeedback were able to learn to adopt the assigned interpretation. The black line indicates the line of best fit. Error bars = ±1 s.e.m., + = p < 0.1

We reasoned that participants with higher decoding accuracy would be receiving higher-fidelity neurofeedback, and consequently they should show a larger effect of neurofeedback on their final interpretation of the story. Consistent with this prediction, there was a significant positive relationship between decoding accuracy and correct interpretation scores (Pearson r = 0.56, p = 0.010; Figure 6b).

### 3.6 Results divided by decoding accuracy

Given that decoding accuracy varied considerably across participants, we assessed if the outcomes described above were different in participants with high vs. low decoding accuracy. For these analyses, we performed a median split for each assigned interpretation group based on decoding accuracy. Thus, 5 participants from the cheating group and 5 participants from the paranoid group were in the “best decoding” group and 5 from each group were in the “worst decoding” group. Next, we recalculated our results within each of these new groups. To combine participants who were assigned different interpretations, we again adjusted scores so that positive values relate to the assigned interpretation and negative values relate to the opposite interpretation.

#### 3.6.1 Story comprehension and interpretation scores

Comprehension scores did not vary significantly between decoding accuracy groups (t(18) = −1.41, p = 0.18), shown in Figure 7a. Interpretation scores, however, were significantly higher for the best compared to worst decoding accuracy group (one-tailed t(18) = 2.44, p = 0.013), which was expected based on the significant positive correlation between decoding accuracy and correct interpretation (Figure 7b). For the 10 participants in the best decoding accuracy group, there was a trend for interpretation scores to have positive (i.e., correct) values, but that trend was not significant (one-tailed t(9) = 1.541, p = 0.079).

**FIGURE 7.**
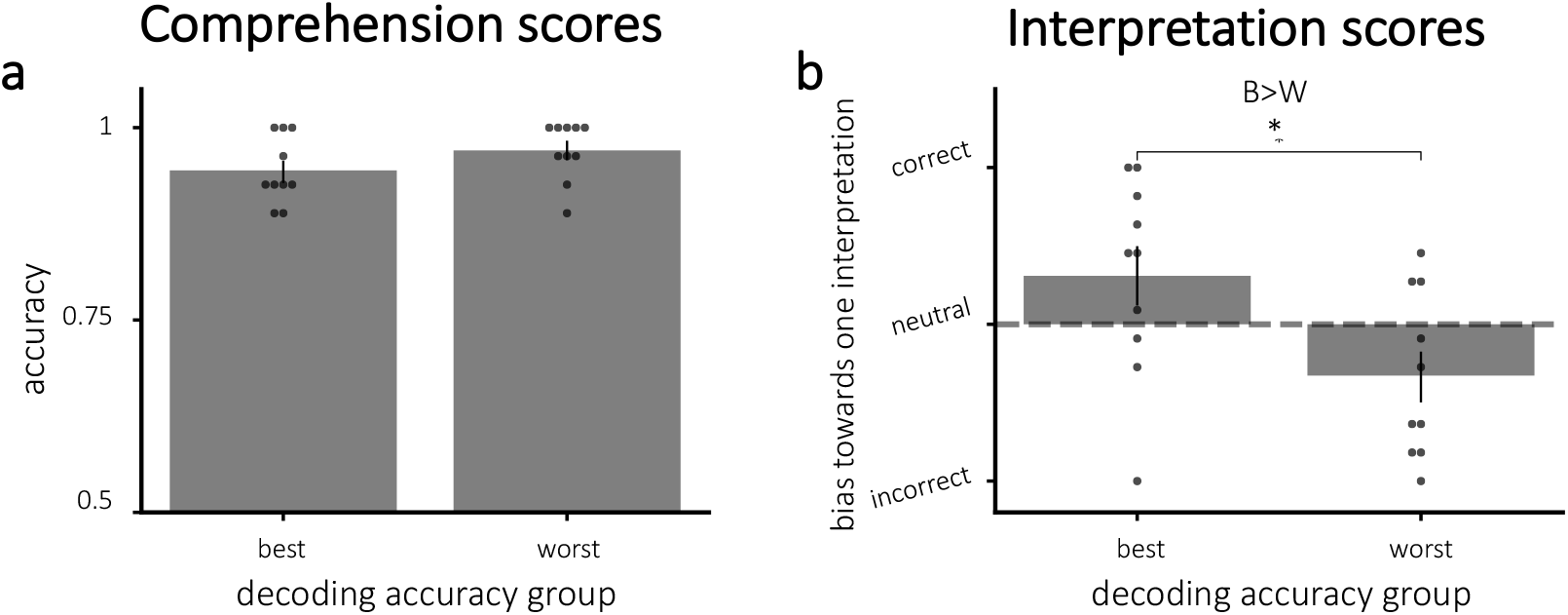
Average scores for **(a)** comprehension and **(b)** interpretation questions, divided by decoding accuracy **(a)** Comprehension scores did not vary significantly between groups. **(b)** Correct interpretation was significantly larger for the participants with more accurate decoding. **Key**: B = best; W = worst. Error bars = ±1 s.e.m. * = p < 0.05

#### 3.6.2 Empathy ratings

The difference in empathy for Arthur and Lee did not vary significantly depending on decoding accuracy (one-tailed t(18) = 0.77, p = 0.23; Figure 8). Still, participants with the best decoding accuracy had a numerically higher mean (indicating empathy scores that fall more in line with the “correct” interpretation) than those with the worst decoding accuracy.

**FIGURE 8.**
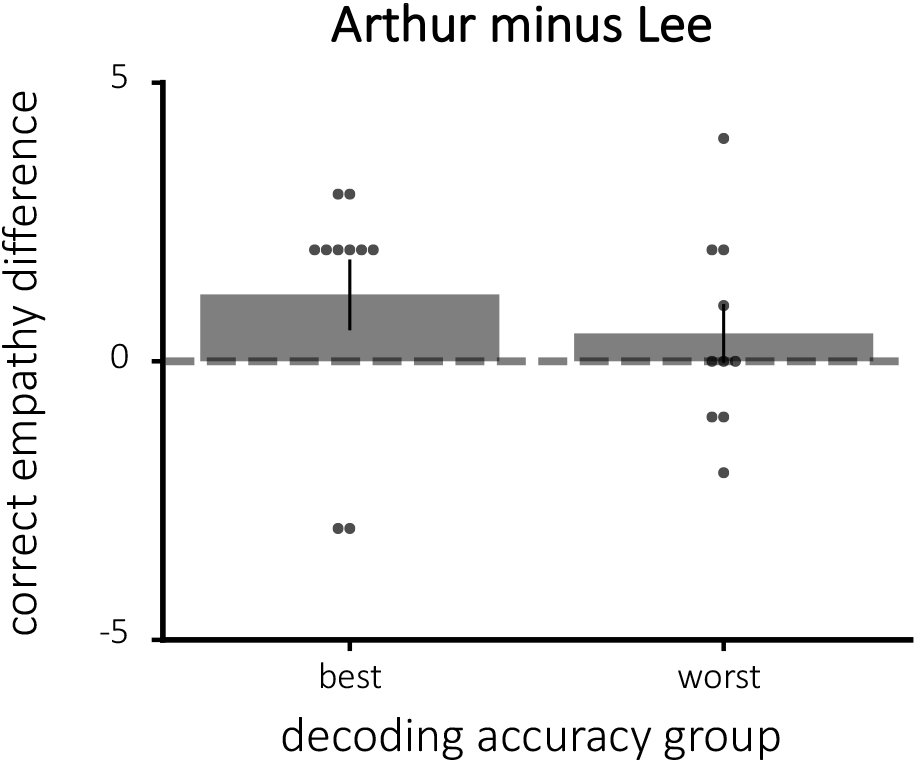
The difference in empathy for Arthur and Lee, divided by decoding accuracy. Here, the empathy difference is adjusted for assigned group such that positive empathy differences indicate the correct direction. There was no significant difference between the best and worst classifier groups in terms of modifying empathy.

#### 3.6.3 Neurofeedback scores

We plotted neurofeedback scores, both in terms of prediction probabilities and normalized neurofeedback scores (Figure 9). Looking at individual stations, we only obtained one significant result after Bonferroni correction for multiple comparisons across stations: For the final station in the first run, neurofeedback scores were higher for participants that had the best decoding accuracy (one-tailed t(18) = 3.40, p = 0.0016 before correction and p < .05 after correction). When averaging over all stations within each run, there was a nonsignificant trend for neurofeedback scores in the fourth run to be higher for participants that had the best decoding accuracy (one-tailed t(17) = 2.36, p = 0.015 before correction; this was no longer significant after Bonferroni correction for four tests).

**FIGURE 9.**
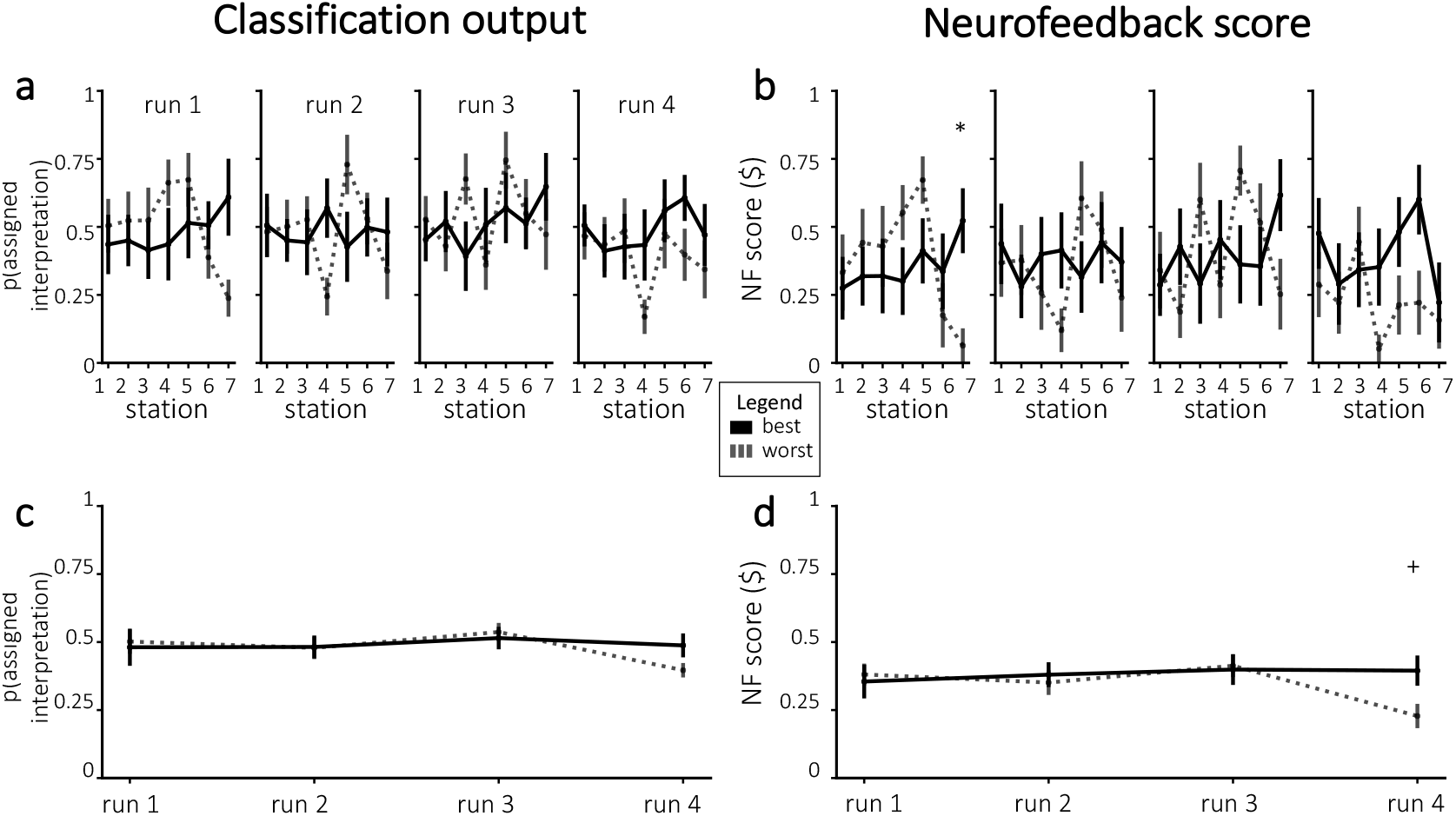
Neurofeedback results, in terms of classification output **(a, c)** and normalized neurofeedback score **(b, d)**. **(a)** Average classifier probability for the assigned interpretation, for the participants with the best and worst decoding accuracy. **(b)** Neurofeedback reward, divided by the participants with the best and worst decoding accuracy. Figures **(c-d)** show the run-wise averages for the same values shown in **(a-b)**, respectively. Error bars = ±1 s.e.m. Significance was corrected for multiple comparisons using Bonferroni correction. * = p < 0.05; + = p< 0.1

#### 3.6.4 Probe responses

As shown in Figure 10, choosing the correct interpretation during probes was closely related to decoding accuracy. For the second-to-last station, participants with the best decoding accuracy were significantly more likely than participants with the worst decoding accuracy to choose the correct assigned interpretation in both run 2 and run 4, after Bonferroni correction for multiple comparisons; there were trending differences between groups for the same station in run 1, and for the preceding two stations in run 4.

**FIGURE 10.**
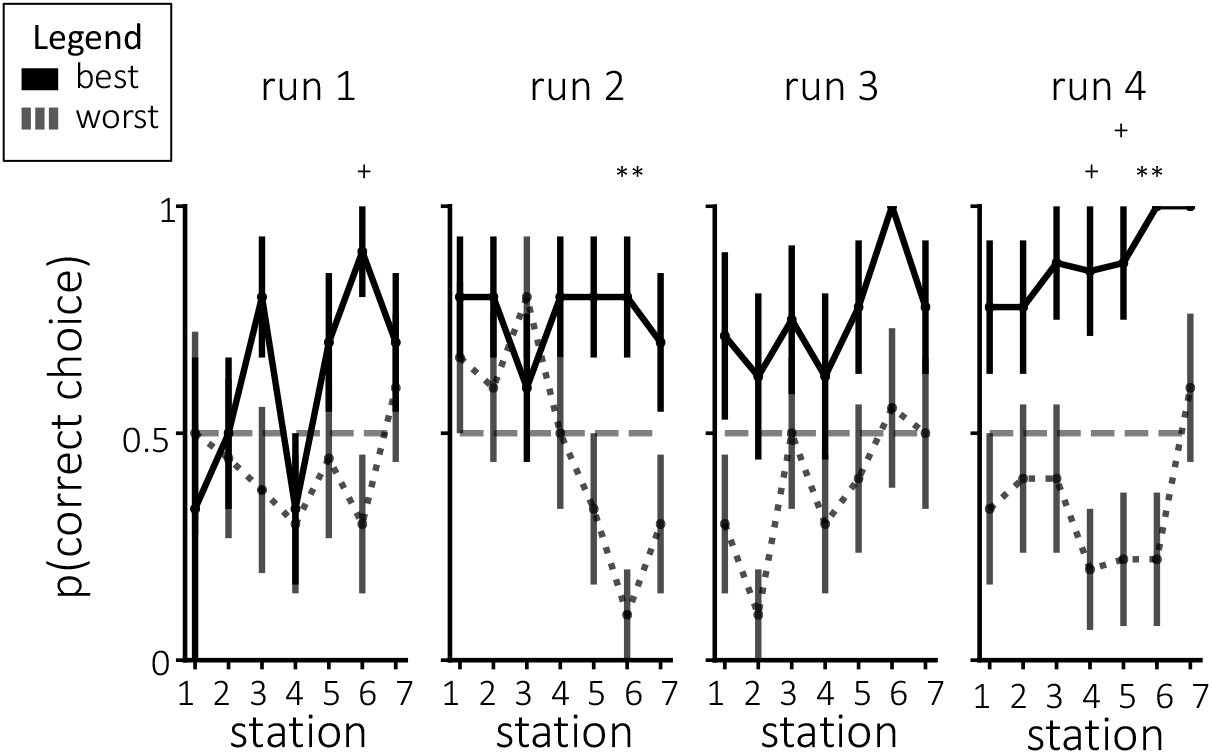
Probe choices split by decoding accuracy. Participants with the most accurate decoding were more likely to choose probes consistent with their assigned interpretation by the end of training. Error bars = ±1 s.e.m. Significance was corrected for multiple comparisons using Bonferroni correction. ** = p < 0.01; + = p < 0.1

## 4 DISCUSSION

In summary, we examined the effect of neurofeedback during a naturalistic spoken story on several behavioral and neural measures of narrative interpretation. Behaviorally, we tracked participants’ overall story interpretation at the end of neurofeedback, their empathy for the different characters at the end of neurofeedback, and also their responses to probes (during neurofeedback) about what interpretation they planned to adopt at each station. Neurally, we used a classifier (trained on previously-collected data from Yeshurun et al., 2017) to track what interpretation participants adopted at each station; we also tracked how much reward participants received, which was tied to how strongly their brain activity matched the assigned interpretation.

When considering the entire participant group, the effects of neurofeedback were fairly weak: Only one of the aforementioned measures (empathy for the characters) showed a statistically reliable effect of neurofeedback. The effects of neurofeedback were somewhat stronger when we grouped participants based on the station-to-station correspondence between their behavioral probe responses and classifier evidence (i.e., was the classifier more likely to favor the “cheating” interpretation at a station when participants behaviorally responded that they planned to adopt the cheating interpretation at that station). This *decoding accuracy* measure shows how well the classifier performs at decoding the interpretation that participants *say* they are adopting, regardless of whether that interpretation is the “correct” (i.e., assigned) interpretation. We found that participants who ranked high on this decoding accuracy measure were more likely to show the predicted effect of neurofeedback on behavioral interpretation scores. When we did a median split based on this measure (dividing participants into “best-decoding” and “worst-decoding” groups), the “best-decoding” participants showed a stronger effect of neurofeedback than the “worst-decoding” participants on behavioral probe responses in run 4 (i.e., they were more likely to indicate that they were adopting the assigned interpretation for upcoming stations); there was a trending (but nonsignificant after correction for multiple comparisons) effect on neurofeedback reward for run 4 (i.e., levels of reward were higher for the “best-decoding” participants, indicating that classifier output was more in line with the assigned interpretation). However, even within this selected group, the effect of neurofeedback on classifier outputs was somewhat weak (the aforementioned trending effect in run 4 reflected participants with the worst decoding accuracy being more strongly nudged to the incorrect interpretation, rather than participants with the best decoding accuracy robustly adopting the correct interpretation).

There are two (non-mutually-exclusive) explanations for why neurofeedback effects were generally larger in the “best-decoding” participants. One possibility is that our classification pipeline did a better job of decoding story interpretations in some participants than others. These participants may have received more accurate (and thus more useful) feedback, leading to larger neurofeedback effects. If some participants failed to show neurofeedback effects because of a poorly functioning classifier, then the most effective way to boost neurofeedback effects in future studies would be to improve our classification pipeline to make it work more consistently across participants (e.g., by trying to improve registration, ROI selection, classifier parameter choices, and the quality of the template data). Another possibility is that the classifier itself was working well in the “worst decoding” participants, but these participants had trouble shifting their interpretations (e.g., a participant might behaviorally signal that they intend to adopt a “cheating” interpretive lens for the upcoming station, but then fail to actually do this). That is, the problem may reflect a cognitive difference across participants (some participants can cleanly shift their interpretations, some cannot) rather than a problem with neural decoding. If this is the case, perhaps screening participants beforehand with various behavioral tasks (e.g., measuring cognitive flexibility) could help exclude participants who will ultimately not benefit from neurofeedback. Based on our current set of results, we can not adjudicate between these two interpretations of why the “worst-decoding” participants did not benefit from neurofeedback.

For the neurofeedback effects that we did observe, we must critically assess the role of *individualized* neurofeedback in driving these effects. The premise of our approach is that measuring a particular individual’s interpretation neurally and using this signature as the basis for feedback is useful for changing their interpretation. Importantly, the presence of a difference between the two neurofeedback groups in our study does not necessarily mean that *individualized neurofeedback* was responsible for this difference. For example, say that everyone strongly adopts the cheating interpretation at a particular point in the story and the classifier registers this. In this scenario, participants in the cheating group will be given “correct” feedback, reinforcing the interpretation, and participants in the paranoid group will be given “incorrect” feedback, leading them to adopt the paranoid interpretation. The key point here is that we could achieve this same effect outside of the scanner (by using behavioral data to pick a point in time when the interpretation is unambiguous, and giving the two groups different feedback at that point). For our study, this scenario is unlikely, because we normalized by the mean interpretation (across all pilot-study participants) when giving feedback - as such, participants adopting the mean interpretation at a particular time point will receive the same (neutral) feedback, regardless of group assignment.

Having said this, more work is needed to establish that individualized feedback is still important. The gold standard for establishing a role for individualized neurofeedback is to include a *yoked control* condition where participants receive feedback based on brain data from another participant in the same condition (deBettencourt et al., 2015; see Sorger et al., 2019, for an overview of different control types for neurofeedback studies). If providing neurofeedback based on another participant “breaks” the neurofeedback effect, this is strong evidence that individualized neurofeedback is important. In our study, this kind of yoked control could be accomplished by replacing the neurofeedback at each station with the neurofeedback that would have been given to another participant from the same overall condition (cheating or paranoid) who had also behaviorally chosen the same interpretation (cheating or paranoid) at that station. Only upon running this control could we then conclude whether or not this rt-fMRI paradigm is able to alter thoughts through individualized neurofeedback.

Returning to the clinical applications discussed at the beginning of the paper: Our interest in this paradigm was driven by the idea that, eventually, we could use this approach to train depressed individuals to interpret realistic scenarios less negatively. What have we learned about the feasibility of this program of research from this study? Overall, we think there are reasons both for optimism and concern. On the optimistic side, we were able to see effects of neurofeedback on both behavioral and neural measures of story interpretation. On the pessimistic side, the effects were weak: The only dependent measure that showed a neurofeedback effect when looking at the entire participant pool was the “empathy” behavioral measure; all of the other effects only showed up when we filtered the participants based on how well the classifier output matched their behavioral probe responses. With the benefit of hindsight, we now think that the “mystery interpretation” approach may not have been the best way to proceed. In the present study, we told participants to explore interpretations and not to lock in on one. We thought that this would help participants pay attention to neurofeedback, but instead we may have encouraged participants to focus too much on hypothesis-testing strategies (e.g., adopting an interpretation contrary to the preferred one, to see if this results in a reduction in reward) - this approach may reduce engagement with the story overall, thereby weakening our effects. Going forward, we think that it might be more effective to simply reveal the assigned interpretation outright, and then use neurofeedback to maximize the degree to which participants adopt and sustain that interpretation over time. This approach would fit better with our long-term clinical goal of using this method to treat depression - there, you would want to inform participants that things will turn out well in the story and then help the depressed participants adopt and sustain that interpretation. Note that, if you reveal the desired interpretation, this makes it less informative to use behavioral interpretation questions as a dependent measure (since participants will know the “right answer”) but neural measures (e.g., the output of a classifier tracking the presence of the correct interpretation) could be used instead.

On a broader level, this work constitutes the first attempt to use rt-fMRI to alter ongoing thoughts related to naturalistic narrative stimuli. In addition to having greater ecological validity than traditional experimental stimuli, naturalistic stimuli have the advantage of producing robust neural responses (Sonkusare et al., 2019; Nastase et al.,2020); they have been shown to be useful for studying individual differences (Vanderwal et al., 2017; Finn et al.,2020; Feilong et al., 2018) and for exposing neural correlates of clinical variables (Rikandi et al., 2017; Finn et al.,2018; Eickhoff et al., 2020; Salmi et al., 2020). Our study highlights both the challenges and future promise of using neurofeedback in a naturalistic context to reshape how we interpret ambiguous situations.

## Acknowledgements

This work was supported by funding from the Intel Corporation to K.A.N., a grant from the John Templeton Foundation to K.A.N., and a grant from the National Institutes of Health (T32MH065214) to A.C.M. The opinions expressed in this publication are those of the authors and do not necessarily reflect those of the funders.

## Conflict of interest

The authors have nothing to disclose.

## Appendices

### A CLASSIFIER TRAINING

In order to provide neurofeedback in real-time, we had to construct an accurate classifier based on previously collected data. In the sections to follow, we discuss our decision-making process for choosing (1) how we preprocessed the training data, and (2) which time points of the story would be used as stations. All analyses and results in this section rely entirely on a previously collected data set (Yeshurun et al., 2017) - they do not use any of the new data that we collected for this paper.

#### A.1 Data acquisition

We used previously-collected data from Yeshurun et al. (2017), where participants were explicitly instructed to adopt one of two different interpretations before listening to the story in the fMRI scanner. This manipulation ensured that the two groups of participants would interpret the story in different ways, allowing the authors to look for neural signatures of interpretation that were shared within each group and differed across groups. The data set included 38 participants: 19 participants who were told that Joanie was cheating on Arthur, and 19 participants who were told that Arthur was paranoid. *Note*: The data published in Yeshurun et al. (2017) included 20 participants per group, but 2 of the participants (one from each group) had start times that did not match the others and were omitted from the analysis for this reason. For more details regarding this data set, see Yeshurun et al. (2017). We preprocessed the neural data from the remaining 38 participants in the same way as detailed above (see Methods, Section 2.4.1), using *fMRIPprep* 1.2.3 (Esteban et al., 2019; Esteban et al., 2018; RRID:SCR_016216).

#### A.2 Finding the optimal classifier

To optimize our classifier, we ran multiple analyses exploring variants of our analysis pipeline. The goal of these analyses was to decide on preprocessing parameters for our neurofeedback analyses and also to decide what time points to use for neurofeedback. All analyses presented in this section are based entirely on the previously-collected data from Yeshurun et al. (2017) - they do not rely on any of the new data collected for this paper.

Figure 11 shows a high-level overview of the classifier pipeline that we used for these parameter-optimization analyses. We discuss each step in detail next.

**FIGURE 11.**
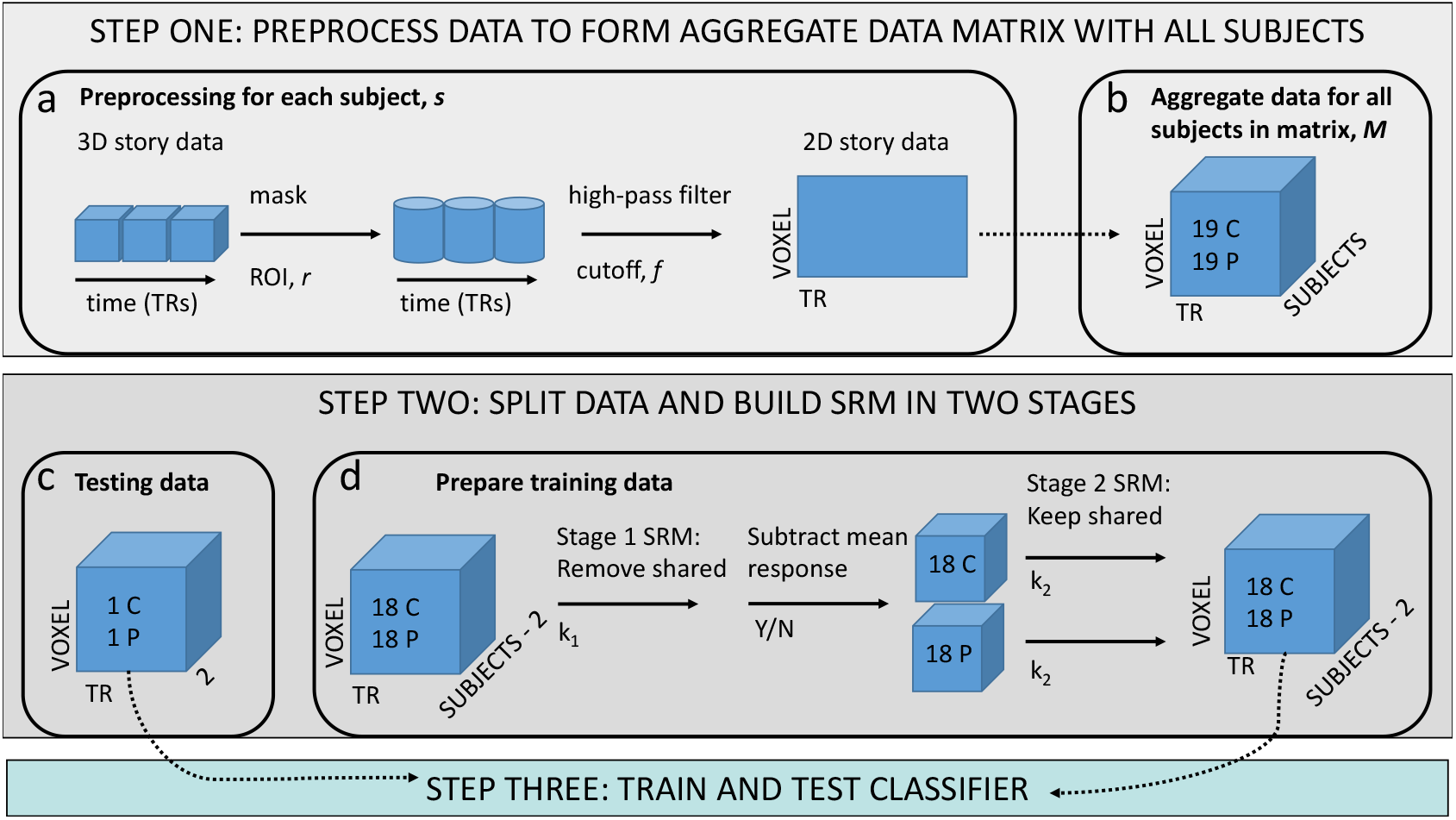
Classifier pipeline for parameter optimization analyses. Before splitting the data into training and test data, we completed the following preprocessing steps for all participants: We masked the story time series given a specific ROI; we high-pass filtered the data based on a particular cutoff (or no cutoff) frequency; and we flattened this time series into a 2D matrix. Next, we randomly selected 2 participants (1 per group) to be left out for testing. For the remaining training data, we randomly selected 18 participants per group (with replacement) so the groups were balanced. Then, we trained a Shared Response Model (SRM) on all training data using k_1_ dimensions and removed the signal that was shared across all participants (regardless of group). Next, we subtracted the mean response (if this step was included). We then trained separate SRMs on each group using k_2_ dimensions and kept only the shared signal within each group. Finally, we trained the classifier on the separate group shared signals (in voxel space). We then used the classifier to predict group labels for the 2 held-out participants. **Key**: C = cheating group; P = paranoid group; SRM = shared response model.

##### A.2.1 Step one: preprocessing

Preprocessing steps (shown in boxes **a-b** of Figure 11) entailed:

1. Removing the first 2 TRs
2. Masking with the given ROI
3. High-pass filtering based on the given cutoff frequency
4. Z-scoring each voxel’s timeseries
5. Combining all participants to form a final matrix

The parameters that were varied for our classifier optimization analysis included:

1. **ROI:** We considered three ROIs for this study. The list of ROIs and the steps taken to create each ROI are shown in Table 1.
2. **High-pass filtering:** Because the story lasted about 12 minutes, we had no a priori knowledge of what filtering cutoff would be optimal in removing noise. Thus, we tested 3 different frequency options for high-pass filtering: (1) no filtering, (2) high-pass filtering with a cutoff of 337.75 s as was done in Yeshurun et al. (2017), and (3) high-pass filtering with a cutoff of 140 s as is commonly done in fMRI studies. These options are represented in Figure 12 as indices 0,1, and 2, respectively.

**TABLE 1.** All ROIs considered in our bootstrap analysis, with a description of how we created each mask. The index on the left corresponds to the ROI key shown in Figure 12.

| Index | Name | Method | N voxels |
| --- | --- | --- | --- |
| 0 | Default Mode Network (DMN) | Yeo et al. (2011):<br>- resampled to functional space | 3,757 voxels |
| 1 | Theory of Mind (TOM) | Neurosynth (Yarkoni et al., 2011):<br>- used the association test only<br>- thresholded at FDR-corrected $p \leq 0.01$ significance<br>- resampled to functional space<br>- masked by average brain mask across all participants | 2,414 voxels |
| 2 | TOM clusters | Yeshurun et al. (2017), Supplementary section (Table 1):<br>- Extracted significant clusters: left TPJ, precuneus, posterior cingulate, vmPFC, dmPFC, rIFG, lIFG, ISTS, and left temporal pole<br>- converted to MNI space<br>- created 10-mm cube around each cluster<br>- resampled to functional space<br>- masked by average brain mask across all participants | 240 voxels |

**FIGURE 12.**
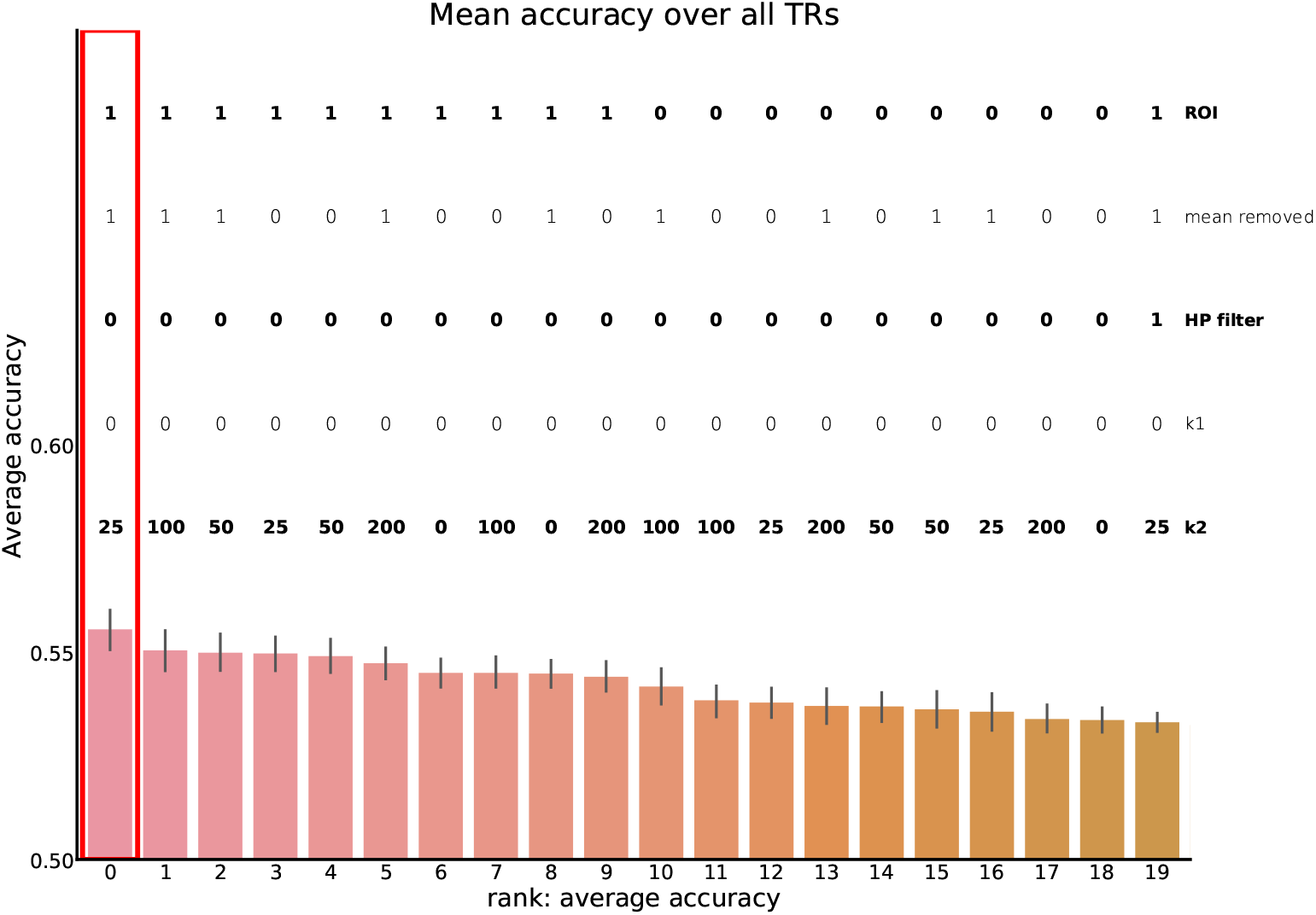
Results from the classifier search. Shown are the top 20 performing parameters for classification ranked by the average accuracy across all TRs. Parameter settings along the x-axis are sorted from best performing (left) to worst performing (right). The red box marks which parameter set was ultimately chosen: ROI = large TOM mask, average signal removed, no high-pass filter, k_1_ = 0 (no first stage SRM), k_2_ = 25. Error bars show 95% CI over bootstrap iterations.

##### A.2.2 Step two: splitting data and cleaning signal with the Shared Response Model (SRM)

We based our SRM approach on Chen et al. (2015). The authors applied a novel two-step shared response model to the data in Yeshurun et al. (2017) in order to improve classification accuracy. Usually, SRM is used to extract the common signal to best represent the stimulus-evoked response to naturalistic stimuli (Chen et al., 2015; Vodrahalli et al., 2018). In this case, because the dataset in Yeshurun et al. (2017) included two groups who listened to the same narrative, we were not interested in the signal that was shared across *all* participants. Rather, we wanted to extract the signal that was common *within each group*, as the shared signals within each of the two groups would likely reflect interpretations. See Chen et al. (2015) for additional mathematical details on SRM analysis and BrainIAK (https://brainiak.org/; Kumar et al., 2020) for details on implementation. We modified their SRM approach slightly for our real-time application, as reported below.

Analysis steps shown in boxes **c-d** of Figure 11 entailed:

1. Randomly selecting two participants (one from each group) to hold out of all subsequent steps and keep as testing data.
2. Randomly resampling 18 participants within each group (with replacement) to build the training data matrix.
3. Training an initial SRM model with k_1_ dimensions using all 38 training participants, and removing the component of the signal that was shared amongst **all training participants**. The purpose of this step was to remove signal pertaining to processing that was common across all participants, regardless of interpretation (e.g., processing relating to low-level sensory features of the stimulus).
4. Subtracting the mean response for each voxel over all participants. This step was done in Yeshurun et al. (2017) - it serves a similar function to running SRM on the full set of participants (i.e., it helps to remove signal that is shared across the two interpretation groups).
5. Separating the groups by assigned interpretation and training another pair of SRMs, one for each interpretation group (separately), using k_2_ dimensions per SRM. This time, we kept only the shared component for each participant. This is intended to highlight shared variance that pertains to the assigned interpretation and remove parts of the signal that are idiosyncratic to particular participants (e.g., thoughts unrelated to the story).
6. Compiling this data into the final training data matrix before testing the classifier. Note that the data were kept in voxel space throughout this analysis pipeline.

The parameters that were varied for our bootstrap analysis included:

1. **k**_1_: We modified the number of dimensions for the first SRM. We also omitted this first SRM (k_1_ = “0”) under one parameter setting.
2. **Subtracting the mean:** We either included this step or not.
3. **k**_2_: We modified the number of dimensions for the two within-group SRMs.

##### A.2.3 Step three: Training and testing

After all preprocessing steps were complete, we then tested the model on the 2 participants that were held out. To obtain confidence intervals for each preprocessing configuration, we ran 1,000 bootstraps of each configuration - for each for these 1,000 bootstraps, we randomly selected one participant per group to be used for testing data, and we resampled from the remaining 18 participants per group (with replacement) to obtain our training set.

We used scikit-learn’s (Pedregosa et al., 2011) SVM classifier (kernel=‘linear’, probability=True) to predict the group label of the 2 left-out participants. To identify which time points were most informative regarding the group label, we trained a separate classifier for each individual TR in the story.

##### A.2.4 Bootstrapping results

The results of our bootstrap analysis are shown in Figure 12. We chose the classifier with the highest accuracy, averaging across all TRs. This classifier corresponded to the following preprocessing steps: ROI = large TOM mask, no high-pass filter, average signal removed, k_1_ =0 (no step 1 SRM), and k_2_ = 25. The performance of this classifier is shown in the red box in Figure 12.

#### A.3 Finding stations

After choosing the classifier with the highest accuracy over all TRs in the story, we wanted to improve the efficacy of neurofeedback by providing feedback at the moments when group interpretations, and thus neural activity, were maximally different. To this end, we used behavioral ratings from Yeshurun et al. (2017) to identify points in the story where interpretations were maximally different. Each of the story’s 179 segments were rated in terms of how much they differed in beliefs, emotions, and intentions depending on the context.

To combine all the ratings into one score, we first z-scored belief, emotion, and intention ratings across raters. Then, we calculated *segment scores* by taking the mean of all ratings at each segment. Thus, larger segment scores meant that those time points were rated as differing more in the story. In Figure 13, the segment scores are indicated by the dashed blue line and the average individual TR accuracy (from our chosen classifier) is plotted in black. The Pearson correlation between the TR accuracy and segment scores was r = 0.21, meaning that the more a segment was rated as differing between contexts, the better the TR classifier performed. This result aligns with similar results reported by Yeshurun et al. (2017) and provides a sanity check, as some parts of the story had nothing to do with the cheating versus paranoid interpretations (such as when the narrator described a chair).

**FIGURE 13.**
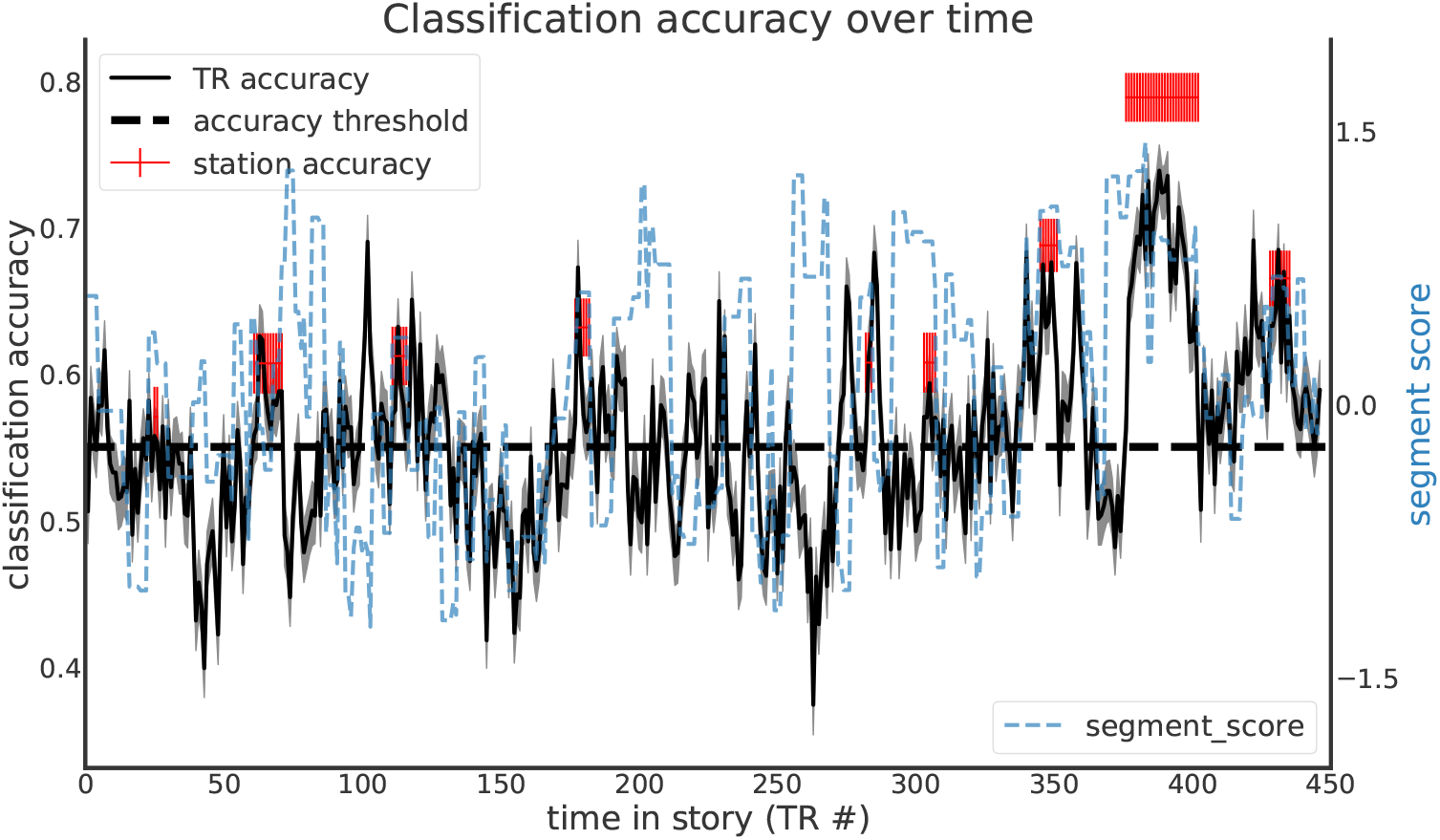
TR and spatiotemporal station accuracy shown over time. The x-axis shows the TR number, while the left y-axis shows the classification accuracy. The **solid black line** represents the individual TR classifier accuracy, with the shaded gray 95% CI over all bootstrap iterations. The **dashed black line** indicates the minimum TR accuracy needed to include a given TR in a station. The **red lines** show chosen stations, with their height indicating classification accuracy, with 95% CI over all bootstrap iterations. The **blue line** shows the segment score ratings (right y-axis). The positive correlation between the TR classification accuracy and segment scores implies that points in the story that were rated to differ in interpretation generally yielded more accurate classification. **NOTE: We did not use the last 2 stations in our main experiment. See Appendix C for details**.

To determine the best time points for neurofeedback, we identified time points meeting the following criteria: (1) individual TR classifier accuracy ≥ 0.55, (2) segment score ≥ 0, and (3) the first 2 conditions satisfied for at least 2 consecutive TRs. We referred to each set of contiguous TRs meeting these criteria as a *station*. After identifying these stations, we extracted a spatiotemporal pattern for each station in each participant by concatenating the spatial response patterns for all time points in the station, resulting in an *n* × *v* vector, where *n* is the number of time points in the station and *v* is the number of voxels in the ROI. We then trained and tested a classifier for each station using the same bootstrapping process that was described above: For each of the 1000 iterations, we randomly sampled participants with replacement to determine the training data and preprocessed the data with the chosen parameters (ROI = large TOM mask, average signal removed, no high-pass filter, k_1_ = 0 (no first stage SRM), k_2_ = 25). The only difference was in stage 3, when we trained and tested a different classifier for each station, instead of each TR.

Out of the stations we tested, we chose final stations that had the highest accuracy and were evenly distributed throughout the story. Overall, the average TR classifier accuracy was 0.56 ± 0.08. The average accuracy for all stations was 0.64 ± 0.1. Thus, incorporating spatiotemporal information allowed us to increase classification accuracy.

### B CLOUD PROCESSING

We designed our real-time pipeline to use a cloud server for processing, and thus minimize dependency on local computing resources (for a similar approach, see Mennen et al., 2021). The cloud-based real-time pipeline was implemented using the RT-Cloud software package (Wallace et al., 2022; Kumar et al., 2020). Once DICOM files arrived at the local Linux machine, they were kept in memory as bytes and immediately sent to the cloud computer. All header information (containing potentially sensitive participant information) was deleted from the DICOM before the file was sent to the cloud. On the cloud server, each BOLD volume was then registered to MNI space, preprocessed, and passed to that station’s classification model for a final neurofeedback score. Finally, a text file was returned to the local Linux machine to update the display. Control of the rt-fMRI pipeline was accessed via a website (requiring security permissions). Figure 14 illustrates both the delineation between local vs. cloud processing, as well as the real-time preprocessing steps. Additionally, Table 2 lists the individual processing steps and software used for each step.

**FIGURE 14.**
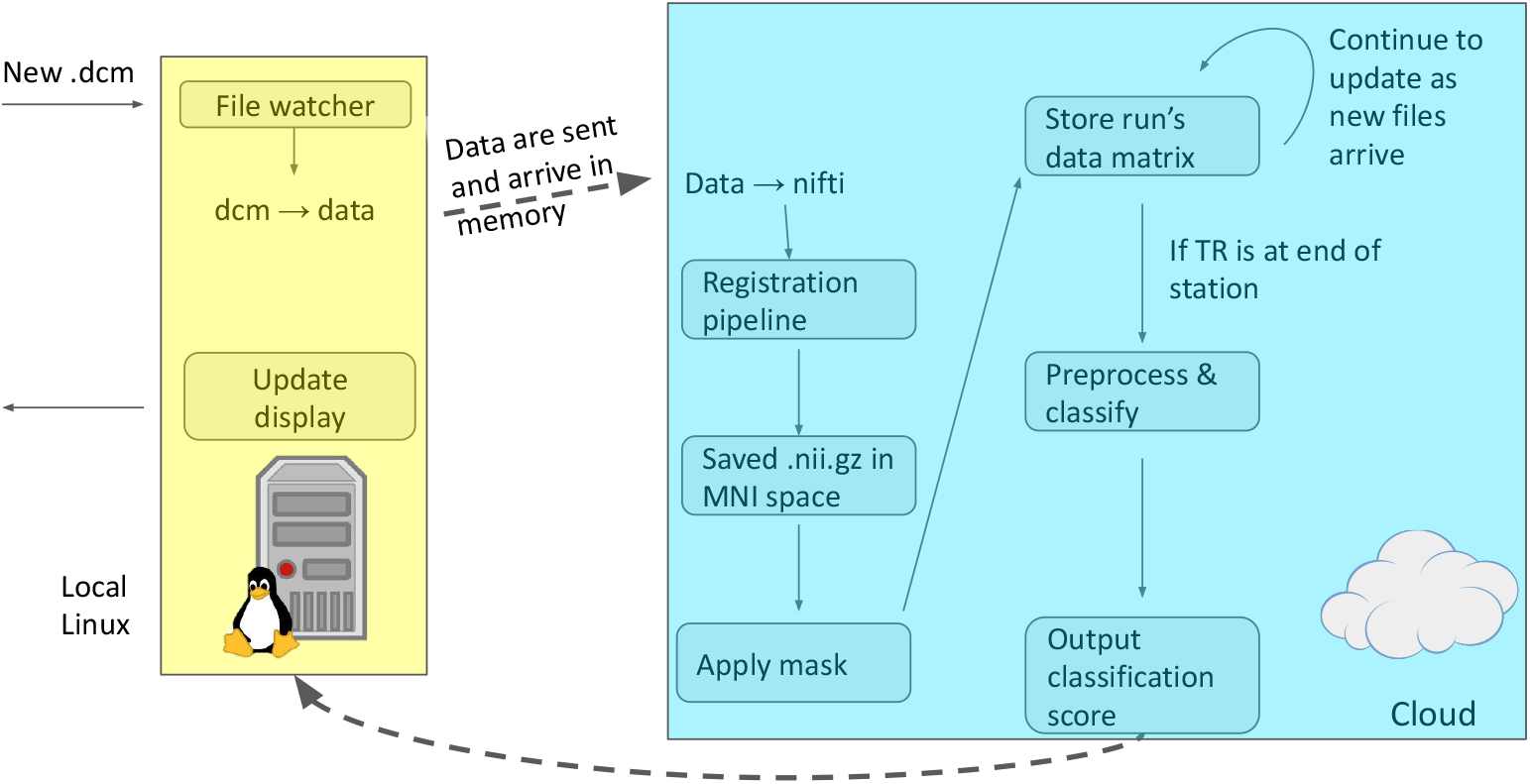
Cloud processing configuration. Data files were sent to the remote cloud server, where they arrived in memory and were converted to a NIfTI file for further processing. A text file containing the neurofeedback score was returned to the local computer.

**TABLE 2.**
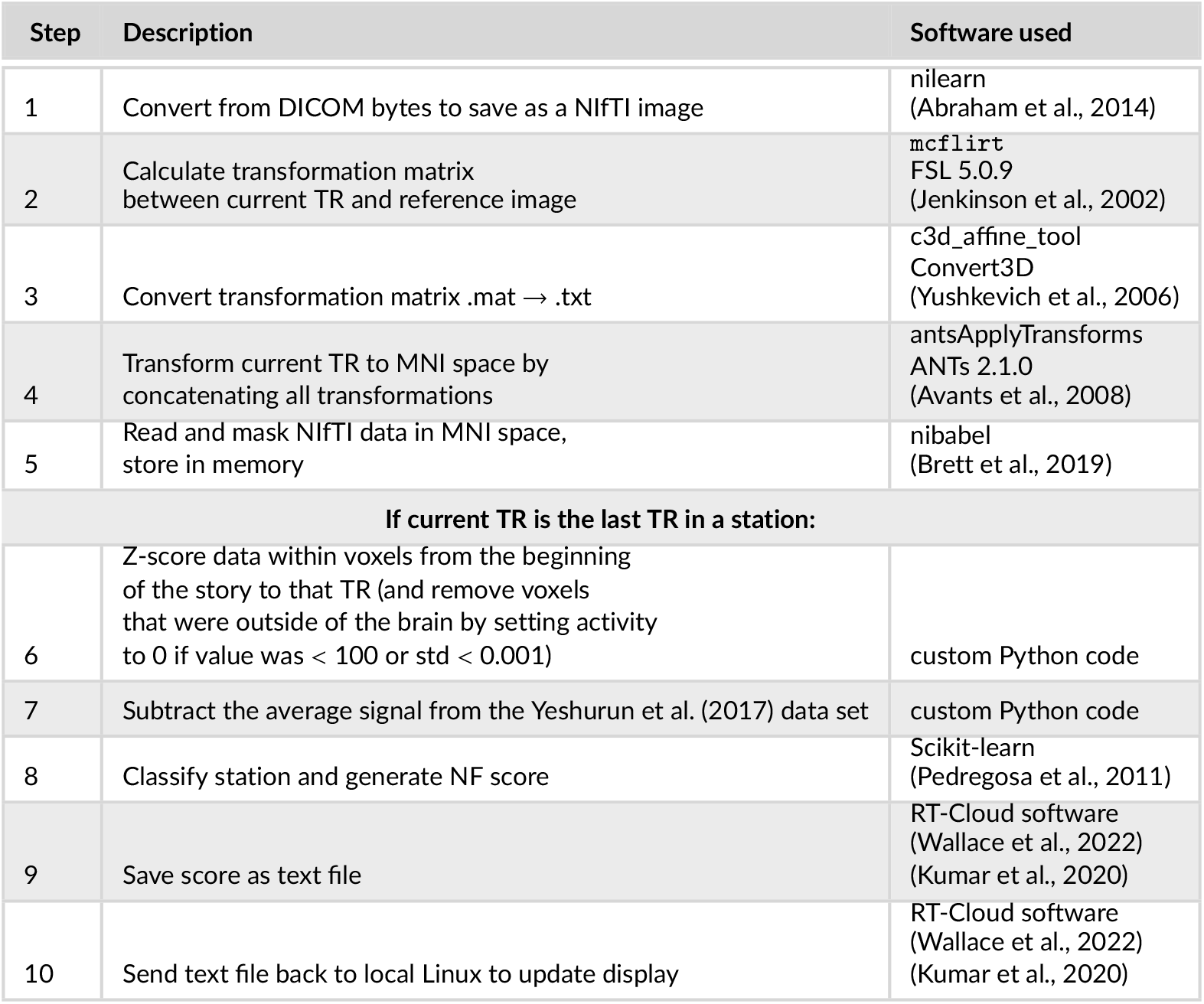
List of preprocessing steps completed on the cloud server for real-time generation of neurofeedback scores.

With regard to timing: As noted earlier, we adjusted for hemodynamic lag by shifting by 3 TRs: If the “station recording” signal was visible to participants from TRs 7 through 12 for a particular station, we analyzed the data from TRs 10 through 15 for that station. Processing steps 1-5 from Table 2 were performed after each TR from the station, and processing steps 6-10 from Table 2 were also performed after the final TR from the station (after TR 15 in this example). We began monitoring for each classification .txt file at the start of the TR immediately following each station (TR 16 in this example) and looked again at the start of the next 9 TRs until the .txt file was received. Considering all participants, runs, and stations, the .txt file was always found at the start of the second TR following the end of the station (TR 17 in this example), with the exception of 2 times when it was received after 6 and 3 TRs due to variations in transmission and processing latency. Once the .txt file was received, the feedback was displayed on the same TR. The upshot of this process was that participants typically had to wait 4 TRs (6 seconds = 3 TR hemodynamic shift, plus one TR for processing) between the offset of the “station recording” signal and the appearance of feedback onscreen.

### C PILOT EXPERIMENT

Before we ran the real-time neurofeedback experiment described in the main text, we piloted an initial version of the experiment. Procedurally, the pilot experiment was the same as the experiment described in the main text, except as noted in the Methods section below. Due to an issue with the SVM classifier, the neurofeedback scores delivered to participants were sometimes incorrect, so the pilot was not a meaningful test of the neurofeedback manipulation. Nonetheless, as described below, we learned some important lessons that we were able to leverage to improve the design of the main experiment.

The most important of these lessons relates to the ambiguity (or lack thereof) of the story. As discussed in the main text, Yeshurun et al. (2017) explicitly told participants which interpretation to adopt ahead of time, and observed neural differences as a function of the assigned interpretation. Our pilot experiment was the first study to explore how participants interpret events in the story stimulus when they are *not* explicitly told which interpretation to adopt ahead of time and (consequently) arrive at interpretations on their own. Naively, we had expected that, given the seeming “ambiguity” of the story, participants’ interpretations would be distributed fairly evenly between the two possibilities (cheating and paranoid) at each time point in the story. What we discovered is that - when participants are not told ahead of time which interpretation to adopt - their interpretations are strongly biased in different directions at different points in the story (i.e., some moments pull participants strongly toward the cheating interpretation, and other moments pull participants strongly toward the paranoid interpretation). Put another way: The ambiguity of the story is not accomplished by making *each moment individually ambiguous*, but rather by see-sawing between moments where different interpretations are more likely, so that when participants *integrate over time* the meaning of the story is ambiguous. Below, we discuss how this discovery led us to re-think our neurofeedback approach, resulting in several changes between the pilot study and the study reported in the main text.

#### C.1 Methods

##### C.1.1 Participants

Nineteen participants consented to participate in this study. Two participants did not understand the task and were excluded from analyses. Data from the remaining 17 participants were retained for further analysis (10 female, 1 left-handed, mean age = 22.4 years). Participants received monetary compensation for their participation, including an additional bonus based on their neurofeedback performance ($20 maximum). The study was approved by Princeton University’s Institutional Review Board.

##### C.1.2 Stimuli

As in our main experiment, participants were randomly assigned to an interpretation group (cheating n = 9, paranoid n = 8). We told all participants about the two possible interpretations before they listened to the story. The participants’ task was to figure out which of the two interpretations was correct through neurofeedback. The auditory narrative itself was identical to the one presented in the main study.

Participants responded to the same comprehension and interpretation questions that were used in the main experiment, with the exception of an additional survey that further probed individual strategies.

##### C.1.3 Procedure

We used the same procedure as described in the main text, except for differences noted below. This time, the mean delay between Visits 1 and 2 was 3.1 days.

The instructions used in the pilot were slightly different from the instructions used in the main experiment. The main-experiment instructions emphasized that the story is purposefully ambiguous with no “correct” interpretation, and that participants should pay attention to the neurofeedback to determine which interpretation we (the experimenters) favored. By contrast, the pilot-experiment instructions presented the task as a mystery, with one interpretation being true. The following points were emphasized in the pilot-experiment instructions:

- Your mission is to figure out the truth behind a phone conversation and solve the mystery
- The neurofeedback scores will reflect how well you’re interpreting the story; a higher score means you are closer to the correct interpretation

The neurofeedback display for the stations (indicating when brain activity was being “recorded”, and then indicating the amount of reward that was accrued from that station) was the same as in the main experiment. However, participants in the pilot experiment were not asked to behaviorally indicate their chosen interpretation before each station (i.e., there were no behavioral responses during the story-listening task).

As in the main experiment, participants listened to the story 4 times with neurofeedback. Note that feedback was provided at the full set of nine stations shown in Figure 13 (the main experiment only used the first seven stations). Once scanning was completed, participants answered the story comprehension and interpretation questions.

##### C.1.4 Data acquisition

All scanning and preprocessing parameters were identical to those described in the main text.

##### C.1.5 Real-time classification

In the main experiment, participants were provided with feedback that was normalized relative to the mean classifier trajectory shown by participants in this pilot study. For example, if the mean classifier score (from the pilot experiment) at a particular time point indicated a .2 probability of the cheating interpretation, and - in the main experiment - a participant in the cheating group showed a showed a classifier probability of .3 for cheating, they received positive feedback, even though (on an absolute scale) the brain state did not favor the cheating interpretation at that time point.

Feedback in the pilot experiment was much simpler: Participants only received positive feedback if the classifier evidence favored the assigned interpretation - that is, participants in the cheating group had to show a classifier probability of cheating > .5 to get positive feedback. Specifically, we used scikit-learn’s (Pedregosa et al., 2011) SVM classifier (kernel=‘linear’, probability=True) and the predict_proba function to convert the output of the SVM classifier to a scalar value *p_c_* indicating the probability of the cheating interpretation.

The neurofeedback score seen by the participant was given by:

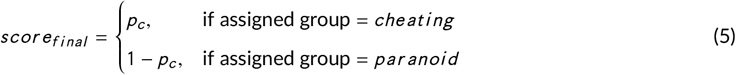

Thus, the higher the neurofeedback score, the more the real-time participant’s neural response matched those of previous participants in the same assigned interpretation group.

Unfortunately, due to a technical problem with the implementation of the predict_proba function, as applied to the SVM classifier in scikit-learn, the neurofeedback scores did not reliably indicate the correct probability. Because of this issue, we do not directly report neurofeedback scores in the results below, nor do we report effects of neurofeedback on participants’ behavioral interpretation scores. Instead, we show results from an offline logistic regression classifier (which did not suffer from this issue with predict_proba) that was applied to the data that we collected during the neurofeedback period. This offline analysis accurately represents the logistic regression classifier’s estimate of participants’ interpretations at each time point in the story.

#### C.2 Results: classifier scores

Figure 15 shows the average classifier-assigned cheating probabilities in the pilot experiment (computed using an offline logistic regression classifier, as described above), split by assigned neurofeedback group. This figure illustrates two important features of the data: First, the two neurofeedback groups showed very similar neural interpretation trajectories. The neurofeedback stations were chosen because they evoked strong neural differences between groups in Yeshurun et al. (2017) when participants were explicitly told which interpretation to take (Figure 13), but the neurofeedback groups did not differ in our study (where participants were not explicitly told which interpretation to take - instead, they had to rely on neurofeedback to guide them). We do not want to overinterpret this, because of the aforementioned bug in the neurofeedback scores, but it is nonetheless striking how similar participants’ trajectories were across groups. Second, and most importantly, the mean classifier probabilities varied sharply across stations, such that many stations were very sharply biased toward particular interpretations. Put another way, when participants were allowed to form their own interpretations (as opposed to being explicitly told which interpretation to use), they were naturally pulled to different interpretations at different points in the story. This was particularly true for the last two stations, where the cheating probability was low for all participants (the second-to-last station corresponds to the part of the story where Arthur tells Lee that Joanie has returned home). For participants in the Yeshurun et al. (2017) study, receiving this information led to stark differences in neural interpretations across the two groups (see Figure 13) - participants in the paranoid group may have felt like this proved their case, whereas participants in the cheating group were forced to conclude that Arthur was an unreliable narrator and he was lying to Lee about Joanie returning home. However, for participants in our study, this new information simply led them to conclude that Arthur was being paranoid.

**FIGURE 15.**
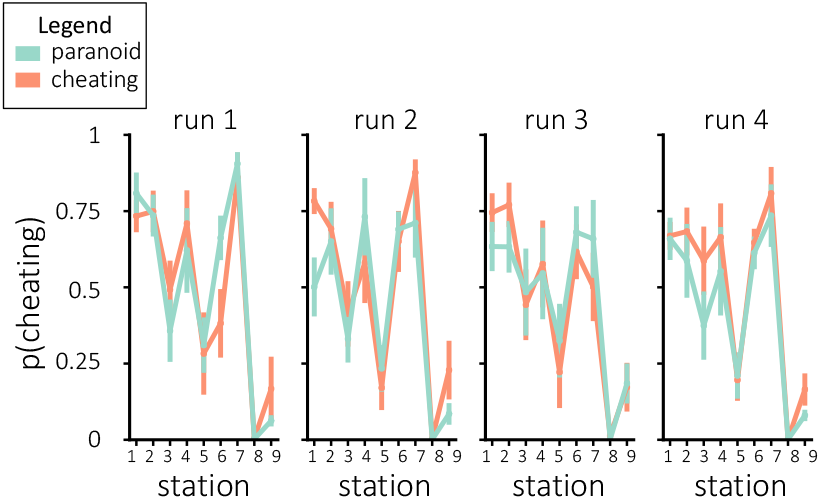
Classifier scores indicating the probability that participants were adopting the cheating interpretation. **These were not the results used in real-time; these results were computed afterward with the same data and a logistic regression classifier.** Error bars represent ±1 s.e.m.

#### C.3 Discussion

Before this pilot, we had assumed from the results in Yeshurun et al. (2017) that the story was uniformly ambiguous. However, the pilot results clearly show that, although the overall story is ambiguous, individual moments in the story were not equally ambiguous. The story’s narrative structure pulls the listener to contradictory interpretations at different points in time.

This lack of “within-timepoint” ambiguity is problematic for our aims, for several reasons. First, if there is too strong of a narrative pull toward one interpretation at a particular time point, this will make it difficult to nudge participants to adopt a contradictory interpretation at that station. Second, lack of within-timepoint ambiguity undermines the purpose of neurofeedback, which is to give participants individualized scores based on their own, varying neural responses to the story. If all participants adopt the same stimulus-driven interpretation at a particular time point, then neural data from a particular participant does not tell you anything new, beyond what you already knew from the neural responses of other participants. For example, based on Figure 15, we can be reasonably sure that a new participant will adopt the paranoid interpretation at station 8, without even measuring their brain activity, so there is no added value to collecting fMRI data at this point.

Based on these results, we opted to make several changes to the paradigm for our own experiment. First, we removed last two neurofeedback stations, where interpretations were strongly biased towards the paranoid interpretation (note that, while we no longer provided neurofeedback at these points in the story, participants still listened to and were scanned during the entire story). However, even after eliminating the most biased stations, there were still some (lesser) biases in interpretation at the other stations. To control for these biases, we decided to provide neurofeedback based on participants’ deviation from the “average neural interpretation trajectory” (where this “average neural interpretation trajectory” was computed by collapsing results across the two conditions in this pilot study) – see the *Classification* section in the main text for details.

We also opted to make changes to the instructions that participants were given. The instructions in the pilot study told participants that there was only one true interpretations, and that they had to discover this one true interpretation. A problem with these instructions was that - once participants decided that they knew the answer - they had no reason to continue attending to the task. To address this problem, we changed the instructions to present the two interpretations as equally probable without one being correct; participants’ task was to discover the interpretation that we wanted them to adopt, not the “true” interpretation. We hoped that this would encourage participants to continue paying attention to the story and to the neurofeedback they received throughout the experiment.

Lastly, for the main experiment, we added behavioral responses during neurofeedback (asking participants, before each station, to tell us which “interpretive lens” they would use for the upcoming station). We thought this would have three benefits: 1) it would promote continued engagement; 2) it would give us a behavioral indicator of what interpretations participants were adopting during the task, which could be used as an additional dependent measure; and 3) it would give us a way of assessing classification accuracy (by comparing the classifier estimate of the participant’s interpretation at a station to the behavioral response at that station).

### D COMPLETE INSTRUCTIONS

We include the instructions that were used in the main experiment in Figure 16.

**FIGURE 16.**
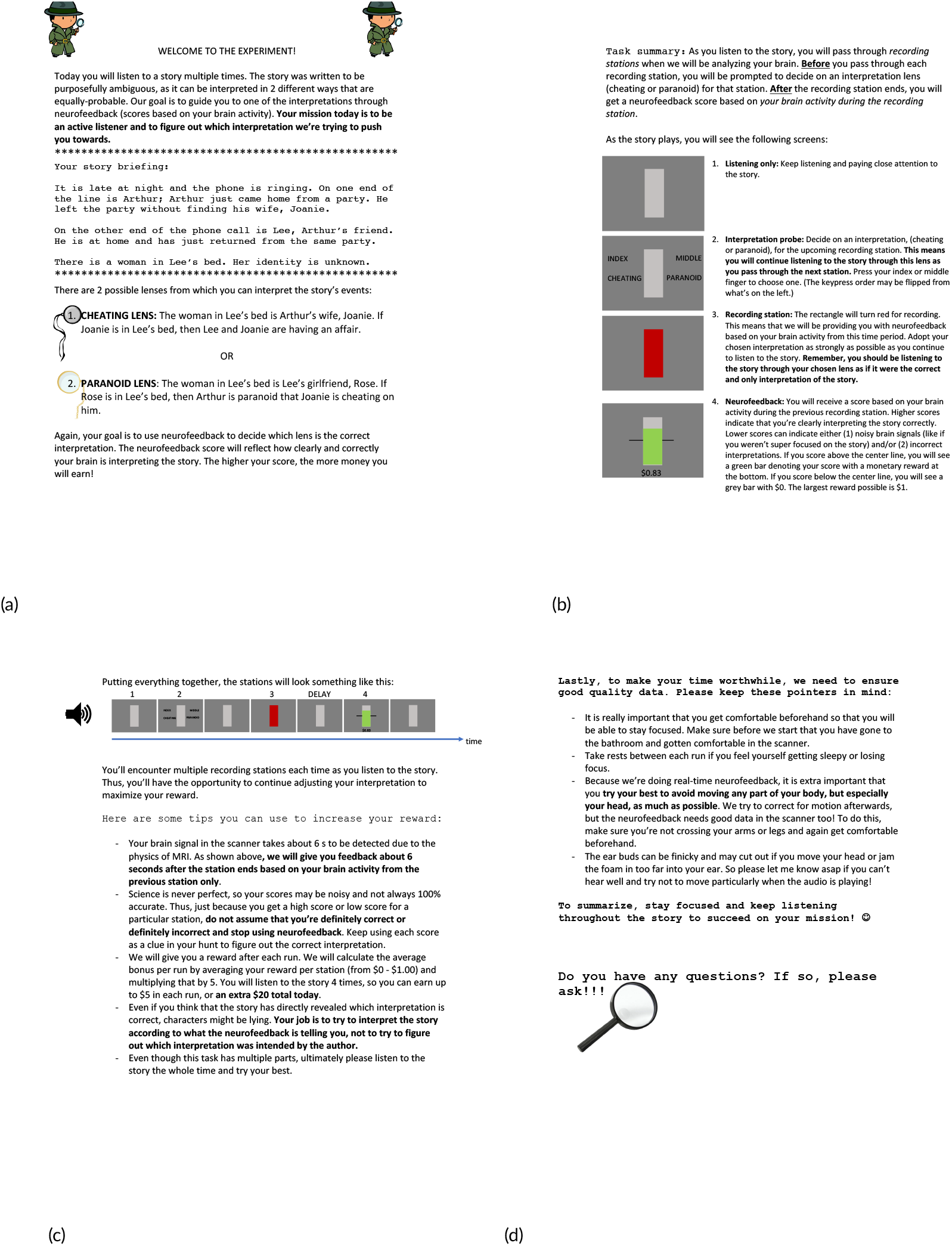
Instructions that span pages 1-4 shown in **a**, **b**, **c**, **d**, respectively.

## Notes

### Competing Interest Statement

The authors have declared no competing interest.

